# Activity-dependent coupling of axonal amphisome trafficking and local norepinephrine signaling *in vivo*

**DOI:** 10.1101/2025.09.23.677999

**Authors:** Ahmed A.A. Aly, Hongbo Jia, Maria Andres-Alonso, Anna Karpova

## Abstract

Amphisomes are autophagy-related organelles formed by the fusion of autophagosomes with late endosomes. In neurons, they contribute to both cargo degradation and signaling, yet their physiological roles *in vivo* remain unclear. Here, we show that axonal amphisome trafficking in highly branched locus coeruleus (*LC*) neurons is regulated by neuronal activity, novelty-associated behavioral state, and axonal norepinephrine (NE) signaling. Using fiber-mediated *in vivo* laser photoconversion and two-photon imaging, combined with chemogenetics and a genetically encoded norepinephrine sensor, we tracked amphisome transport in the intact brains. We show that organelles generated within distal *LC* axons projecting to the prefrontal and motor cortices (*PFC/M1*) undergo retrograde trafficking across the entire axonal length toward somatic compartments. Amphisome trafficking dynamics are bidirectionally regulated by presynaptic adrenergic signaling through opposing cAMP/PKA-dependent mechanisms. *In vivo,* elevated local autoreceptor-mediated norepinephrine signaling, involving association of activated β2-adrenergic autoreceptors with SIPA1L2-positive amphisomes, constrains processive retrograde trafficking by promoting transient immobilization and localized, non-directional “jittery” motility states. Conversely, reduced local norepinephrine signaling promotes amphisome mobilization, and chemogenetic engagement of Gi-coupled signaling accelerated transport by reducing transient immobilization events and reinforcing directional processivity, thereby facilitating cargo delivery toward somatic degradative compartments. Together, these findings identify a local neuromodulatory mechanism linking norepinephrine signaling to axonal amphisome trafficking *in vivo* and suggest that neuromodulatory states, such as novelty- and wakefulness-associated *LC* activity, as well as sleep-associated silencing of *LC* activity, regulate neuronal proteostasis through local control of autophagic cargo transport.

## INTRODUCTION

Maintaining proteostasis across the extraordinary length of axons, which undergo extensive membrane turnover associated with sustained neurotransmission, poses a fundamental challenge for neurons. Autophagy is critical to this process, particularly in distal axons, where autophagosomes are continuously generated and undergo dynein-mediated transport over long distances to the soma, where lysosomes are enriched (Lee et al., 2011; Maday et al., 2012; Maday and Holzbaur, 2014; Cason et al., 2021; Sidibe et al., 2022; Lie et al., 2022). Neuronal amphisomes, hybrid organelles formed by the fusion of autophagosomes with late endosomes, represent key intermediates in this pathway (Wijdeven et al., 2016; Andres-Alonso et al., 2019; Karpova et al., 2025). These organelles are long-lived in neurons and undergo processive retrograde transport over long axonal distances. Their motility and processivity are regulated by a scaffold-motor regulatory network involving RapGTPase-activating protein SIPA1L2 and dynein adaptor Snapin, which couples autophagic and late endosomal compartments to retrograde transport machinery (Cai et al., 2010; Zhou et al., 2012; Cheng et al., 2015; Di Giovanni and Sheng, 2015; Andres-Alonso et al., 2019). This regulation enables diverse motility states of autophagic vesicles, including processive transport, stationary pauses, and prolonged dwell states with variable durations (Jahreiss et al., 2008; Kimura et al., 2008; Cai et al., 2010; Maday and Holzbaur, 2016; Wijdeven et al., 2016; Andres-Alonso et al., 2019; Cason et al., 2021), emphasizing local mechanisms of transport control that may critically regulate the delivery of autophagic cargo to somatic degradative compartments. Beyond their degradative function, accumulating evidence indicates that neuronal amphisomes act as intracellular signaling platforms that regulate synaptic vesicle release and support local protein synthesis (Kononenko et al., 2017; Andres-Alonso et al., 2019, 2026; Karpova et al., 2025). In fact, BDNF/TrkB signaling from amphisomes is essential for BDNF-dependent gene expression in neurons (Kononenko et al., 2017), and the local regulation of TrkB-signaling at synaptic boutons by SIPA1L2 enhances neurotransmission, thereby linking autophagy-related organelles to synaptic signaling pathways (Andres-Alonso et al., 2019). More recent work further suggests that amphisome formation and TrkB signaling at synaptic boutons support local synthesis of presynaptic proteins, indicating a dual role of amphisomes in protein degradation and synthesis (Andres-Alonso et al., 2026). Little is known, however, about the *in vivo* dynamics and circuit-specific functions of amphisomes, and virtually nothing is known about how their trafficking and signaling in long-range projecting neurons are regulated by neuronal activity and autoreceptor activation. Consequently, two fundamental questions remain unresolved: how amphisomes ensure efficient cargo delivery to somatic degradative compartments within defined neuronal circuits and whether they also mediate neuromodulatory signaling in vivo through autoreceptor activation.

*Locus coeruleus* norepinephrine-releasing neurons provide an attractive system for addressing these questions. Despite their small number (fewer than ∼3,000 neurons in rodents (Sara, 2009) and ∼50,000 in humans (Mouton et al., 1994)), *LC* neurons exert widespread modulatory control over arousal, attention, stress responses, and learning through highly branched axonal projections that innervate large portions of the forebrain (Samuels et al., 2008; Sara, 2009; Wagatsuma et al., 2018; Suárez-Pereira et al., 2022; reviewed in Berridge & Waterhouse, 2003; Sara & Bouret, 2012; Chandler et al., 2019; Ross & Van Bockstaele, 2021; Breton-Provencher and Sur, 2019). Their activity is tightly coupled to behavioral state and sleep-wake dynamics, shaping the network activity and memory processes (Kjaerby et al., 2022). Morphologically, *LC* neurons possess highly extended and densely branched axonal arbors (Descarries et al., 1977) that communicate via local volume transmission (McKinney et al., 2023; Chandler et al., 2019; Fuxe et al., 2015; Toyoda et al., 2022), placing substantial demands on axonal proteostasis and autophagic cargo trafficking (Agster et al., 2013; Schwarz & Luo, 2015). These features, together with the accessibility of *LC* axons projecting to the medial prefrontal cortex and motor cortex (*M1*) for *in vivo* imaging, render this pathway a powerful model system for studying the regulation of amphisome trafficking and signaling in the intact mouse brain.

## RESULTS

### SIPA1L2/LC3-positive amphisomes exist in *LC* axons projecting to the prefrontal cortex

In cultured neurons, autophagic vesicles form in distal axons and undergo long-range retrograde transport to the soma for lysosomal degradation (Maday et al., 2012). In a previous work, we showed that trafficking and signaling of TrkB-containing amphisomes in axons of pyramidal neurons is tightly regulated by the RapGAP SIPA1L2 (Andres-Alonso et al., 2019). Dynein-snapin complexes mediate the retrograde organelle transport where binding of LC3 to SIPA1L2 enhances its RapGAP activity, thereby attenuating TrkB-induced Rap signaling and slowing organelle transport (**Fig. 1A**). Under conditions of high neurotransmission, PKA-dependent phosphorylation of snapin and SIPA1L2 immobilizes amphisomes and it allows TrkB signaling by terminating SIPA1L2’s RapGAP activity, facilitating glutamate release (**Fig. 1A**). First, to determine whether amphisomes exist *in vivo* in distal *LC* axons, we unilaterally injected a Cre-inducible adeno-associated virus (AAV) expressing GFP as a cytoplasmic fill into the *LC* of dopamine β-hydroxylase Cre mice (Dbh-Cre; B6.Cg-Dbh^tm3.2(cre)Pjen/J; Tillage et al., 2020). Cre recombinase expression from the endogenous *Dbh* locus enabled selective labeling of *LC* neurons and visualization of their distal projections (**Fig. S1A-B**). Consistent with previous work, immunohistochemical analysis confirmed that GFP-labelled *LC* neurons were indeed positive for LC3B and SIPA1L2, with both proteins co-localized in GFP-labelled *LC* axonal varicosities resembling synaptic boutons (**Fig. 1B, C; Figure S1C**). LC3B puncta also co-localized with snapin at these sites (**Fig. S1D, E**). Finally, the hybrid identity of these vesicles as amphisomes was further supported by immunodetection of the late endosome marker Rab7 near LC3/SIPA1L2 clusters (**Fig. S1F, G**). To examine their mobility in distal *LC* axons *in vivo*, we used a Cre-dependent, *LC*-specific labelling strategy to express mNeonGreen-LC3 (AAV-EF1α-DIO-mNeonGreen-LC3B) and performed two-photon imaging through a transcranial window over *PFC/M1*, a canonical *LC* target (Sara, 2009; Chandler et al., 2014; Totah et al., 2018; **Fig. 1D; Fig. S1H, I**). Time-lapse imaging revealed that amphisomes exhibit robust motility along axons during steady *LC* activity, possibly mediated by SIPA1L2 and its interaction with the dynein adaptor snapin (Andres-Alonso et al., 2019, **Fig. 1D, E**). Quantitative analysis of amphisome trajectories *in vivo*, based on velocity and displacement ratio (total travelled distance over displacement distance) during steady *LC* activity, revealed pronounced heterogeneity in both speed and motility patterns (**Fig. 1F-H**). Some amphisomes moved unidirectionally at ∼0.1 µm/s, whereas others displayed jittery trajectories with fluctuating velocities (**Fig. 1F-H, S1J**). To determine whether these distinct trafficking behaviours might reflect functionally specialized organelle subpopulations, we performed principal component analysis (PCA) using average velocity, maximum velocity, and displacement ratio as quantitative descriptors of amphisome motility. PCA resolved three major populations of amphisomes based on their trafficking dynamics: slow unidirectional, slow jittery, and fast jittery trajectories (**Fig. 1I and S1J**). Collectively, these findings reveal substantial heterogeneity in the motility states and trafficking dynamics of amphisomes within distal *LC* axons projecting to the *PFC/M1 in vivo* under steady-state *LC* activity.

**Figure 1.**
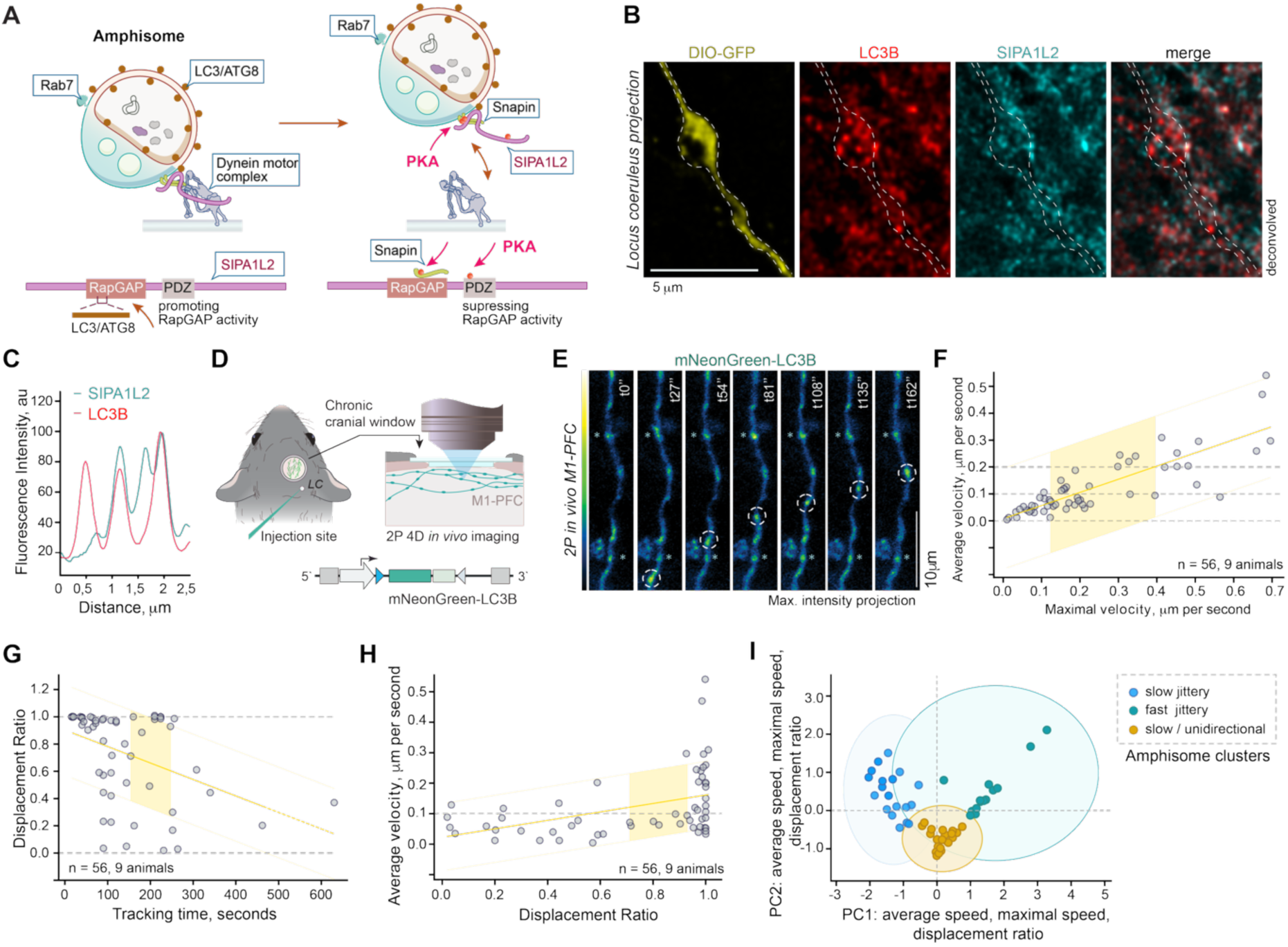
SIPA1L2/LC3B-positive amphisomes localize to *LC* axons projecting to the *PFC/M1* and exhibit three distinct motility patterns. **(A)**. Neuronal amphisomes are hybrid organelles on the autophagy pathway that combine degradative and signalling functions as they travel along long-distance projecting axons. The Rap GTPase-activating protein SIPA1L2 binds LC3/ATG8 and Snapin, thereby linking amphisomes to the dynein motor. Through this interaction, SIPA1L2 governs the speed of retrograde trafficking and modulates local signalling at synaptic boutons (Andres-Alonso et al., 2019, Karpova et al., 2025). **(B)** Confocal image showing a virally expressed GFP-labelled *LC* axon with endogenous SIPA1L2 and LC3B immunoreactivity within an axonal varicosity resembling a synaptic bouton, and **(C)** the corresponding intensity profile. **(D)**. Schematic of *in vivo* two-photon imaging of mNeonGreen-LC3B-labeled amphisomes in *LC* axons projecting to the *PFC/M1*. **(E)**. Representative *in vivo* time-lapse frames acquired through a transcranial window over the *PFC/M1*. Dashed circles highlight amphisomes moving along *LC* axons, whereas asterisks indicate mNeonGreen-LC3B puncta that remained immobile during the image acquisition. **(F)**. Linear correlation between the average and maximum velocities of LC3B-positive amphisomes, reflecting the distance each vesicle travels over time. **(G)**. Correlation between the total track duration and the displacement ratio of LC3B-positive amphisomes. **(H)**. Correlation between displacement ratio and average velocity, illustrating the heterogeneity of amphisome behaviour during spontaneous *LC* activity. **(I)**. PCA of three key features identifies three principal clusters of LC3B-positive amphisomes, each exhibiting distinct trafficking patterns: slow unidirectional, slow jittery, and fast jittery.

### Amphisomes originating in distal axons traverse the entire distance to reach the *LC* soma *in vivo*

Distal axons are generally considered to have limited local degradative capacity, as previously shown in cortical regions (Lie et al., 2021), thereby requiring retrograde transport of autophagic cargo to the soma. Given the observed heterogeneity in amphisome motility states, we therefore quantified lysosomal machinery in distal *LC* projections to the *PFC/M1* to assess whether a subset of vesicles may undergo local fusion with degradative organelles. Axons were labelled with cytoplasmic GFP (**Fig. S1A**), followed by immunohistochemical detection of the lysosomal marker LAMP2A and three-dimensional GFP isosurface masking to restrict quantification to *LC* axonal compartments (**Fig. 2A, B; Fig. S2A**). Quantitative analysis revealed that LAMP2A-positive puncta were only sporadically detected within distal axons, at negligible frequencies compared to the surrounding cellular environment (**Fig. 2A-C and S2A**). This suggests that amphisomes likely remain largely stable during their retrograde transport in *LC* axons due to the limited access to lysosomal compartments in *LC* axons. To determine whether distal amphisomes can traverse the full axonal length to *LC* somata *in vivo*, we used a photoconversion-based labelling strategy. Specifically, in Dbh-Cre mice, *LC* amphisomes were labelled with the photoconvertible fluorophore mEOS4 fused to LC3B (AAV-EF1α-DIO-mEOS4-LC3B; **Fig. 2D**), and an optical fiber was implanted in the *mPFC* to allow local photoconversion (**Fig. 2D; Fig. S2B, C**), as previously described (Yizhar et al., 2011). Subsequent analysis revealed significantly elevated photoconverted “red” puncta in *LC* somata compared with non-stimulated controls (**Fig. 2E**; **Fig. S2B-D**). mEOS4-LC3B-puncta were observed at axonal varicosities (**Fig. S2E**) and, finally, multiple photoconverted puncta were detected in TH-positive neurites adjacent to *LC* somata within 5-6 hours, confirming that distal amphisomes reach neuronal somata *in vivo* (**Fig. 2F**). Colocalization with SIPA1L2 further supported the identification of these puncta as amphisomes (**Fig. 2G, H**).

**Figure 2.**
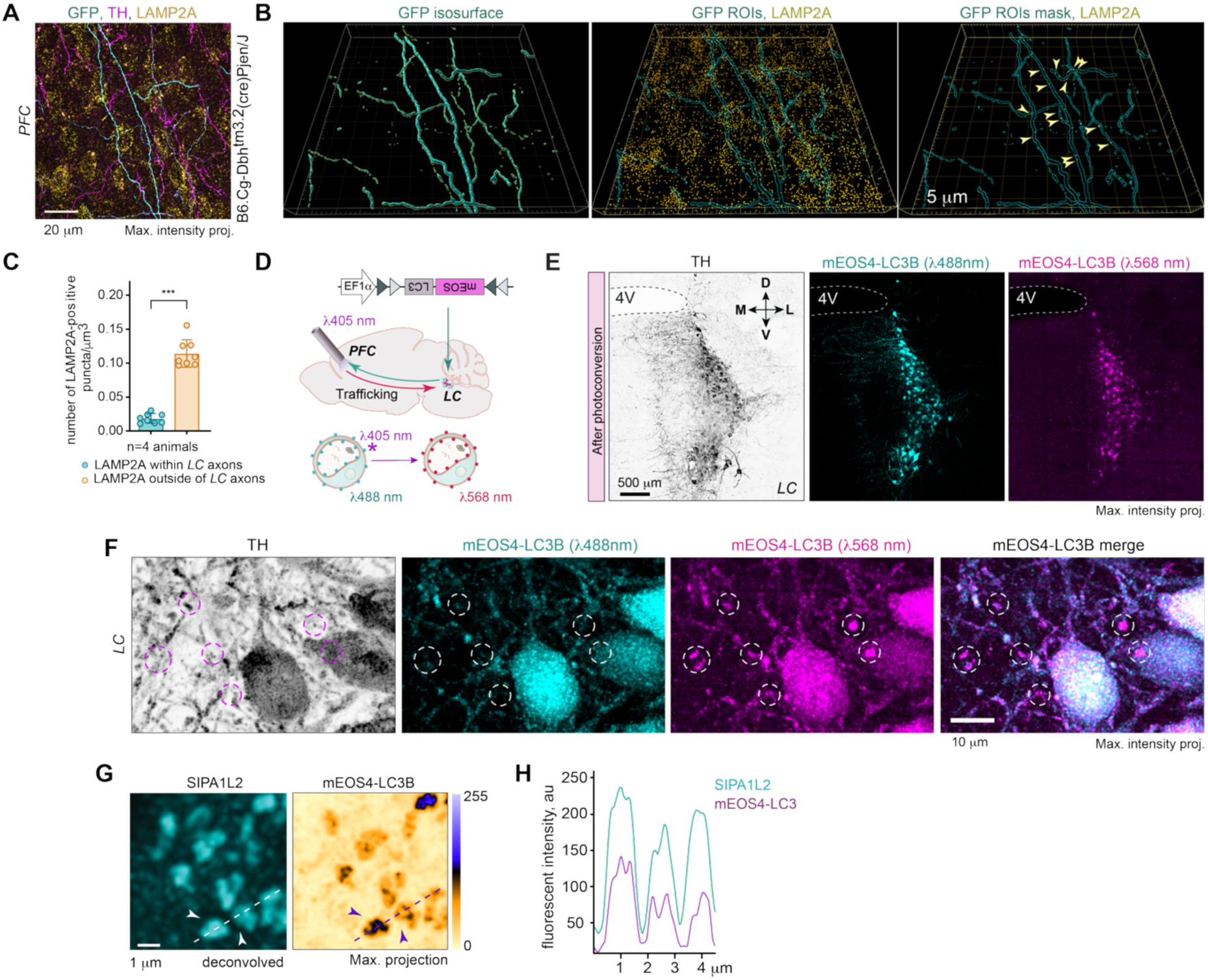
LAMP2A-positive lysosomes are sparsely distributed in *LC* axons projecting to the *PFC/M1*, and amphisomes originating in distal axons traverse long distances to the *LC* soma. **(A)**. Representative confocal maximum-intensity projection of GFP-labelled *LC* axons in the *PFC* co-immunolabeled with anti-TH and anti-LAMP2A antibodies. **(B)**. Three-dimensional reconstructions generated with Imaris (*Bitplane*) software. Left: 3D isosurface rendering of GFP-labelled *LC* axonal projections, illustrating the dense *LC* network in the *PFC*. Middle: Colocalization of GFP-labelled *LC* axons (cyan) with LAMP2a immunoreactivity (yellow). Right: LAMP2A immunoreactivity masked by GFP-defined ROIs (cyan). Box size: 5 µm. **(C)**. Quantification of LAMP2A-positive puncta within GFP-labelled LC axons compared with the surrounding PFC neuropil. Data are presented as mean ± SEM; ***p < 0.001, two-tailed *t-test*. **(D)**. Schematic of the experimental strategy for *in vivo* photoconversion. **(E)**. Confocal tile-scan depicting photoconverted mEOS4-LC3B fluorescence in *LC* neurons following distal photoconversion in the *PFC*. Anti-TH immunostaining labels *LC* neurons. **(F)**. Representative confocal image showing photoconverted mEOS4-LC3B puncta in the vicinity of *LC* somata. **(G)**. Confocal images of *LC* neurons expressing mEOS4-LC3B co-labelled with antibodies against SIPA1L2. **(H)**. Line-intensity profile demonstrating colocalization of somatic mEOS4-LC3B and SIPA1L2.

### Localized stopovers and jittery amphisome dynamics are associated with enhanced NE release, define the axonal NE microenvironment, and shape axonal-to-somatic amphisome delivery

In primary hippocampal neurons, amphisome dynamics are regulated by cAMP/PKA-dependent mechanisms involving the RapGAP SIPA1L2 and the dynein adaptor Snapin, both of which are PKA substrates implicated in amphisome immobilization (Zhou et al., 2012; Cheng et al., 2015; Andres-Alonso et al., 2019). In contrast, inactivation of SIPA1L2 permits TrkB/Rap1 signalling to facilitate synaptic vesicle release (**Fig. 1A).** We first hypothesized that amphisome stopover events in *LC* axons projecting to *PFC/M1,* restricting processive retrograde delivery of autophagic cargo toward proximal axonal regions and the soma, are associated with local spontaneous NE signalling that differentially engages Gs-coupled β-adrenergic receptors and Gi-coupled α2-adrenergic autoreceptors, thereby bidirectionally regulating cAMP/PKA signalling (Strosberg, 1993). We further posited that motility-restricted amphisomes in turn, shape the local axonal NE microenvironment, consistent with a signalling role for amphisomes (Andres-Alonso et al., 2019). To test the relationship between local NE and amphisome dynamics, we combined the genetically encoded high-affinity NE sensor GRAB_NE2h, which enables rapid and selective *in vivo* detection of axonal NE dynamics (Feng et al., 2019, 2024), with simultaneous two-photon *in vivo* imaging of mRuby3-LC3B-labeled amphisomes in *LC* axons (**Fig. 3A-B and S3A-C**). First, spatiotemporal analysis of amphisome stopovers in *LC* axons, performed using the TrackMate (Tinevez et al., 2017; Simon Youssef et al., 2011), revealed that immobilization events coincided with increases in GRAB_NE2h fluorescence that persisted for at least two seconds following amphisome transient immobilization (**Fig. 3B-F**). Intriguingly, correlation analysis revealed that prolonged amphisome immobilization coincided with sustained local elevations in GRAB_NE2h fluorescence, suggesting that immobilized amphisomes act as focal sites that shape the local norepinephrine signalling microenvironment **(Fig. 3G)**. Consistent with cAMP/PKA-dependent regulation of axonal motor-cargo coupling and transport state transitions, including control of dynein processivity and pausing behaviour (Cai et al., 2010; Zhou et al., 2012; Maday et al., 2012; Maday & Holzbaur, 2014), we reasoned that localized “jittery” amphisome motility states, which further restrict processive retrograde progression of amphisomes, may also be associated with local axonal norepinephrine dynamics. Intriguingly, we found that transitions into the “jittery” motility state preceded the onset of elevated local axonal norepinephrine levels, which remained persistently elevated for at least 4 seconds *in vivo* **(Fig. 3H-K)**. Persistent norepinephrine elevation throughout this confined motility state further supports a role for amphisomes in shaping the local norepinephrine signalling microenvironment.

**Figure 3.**
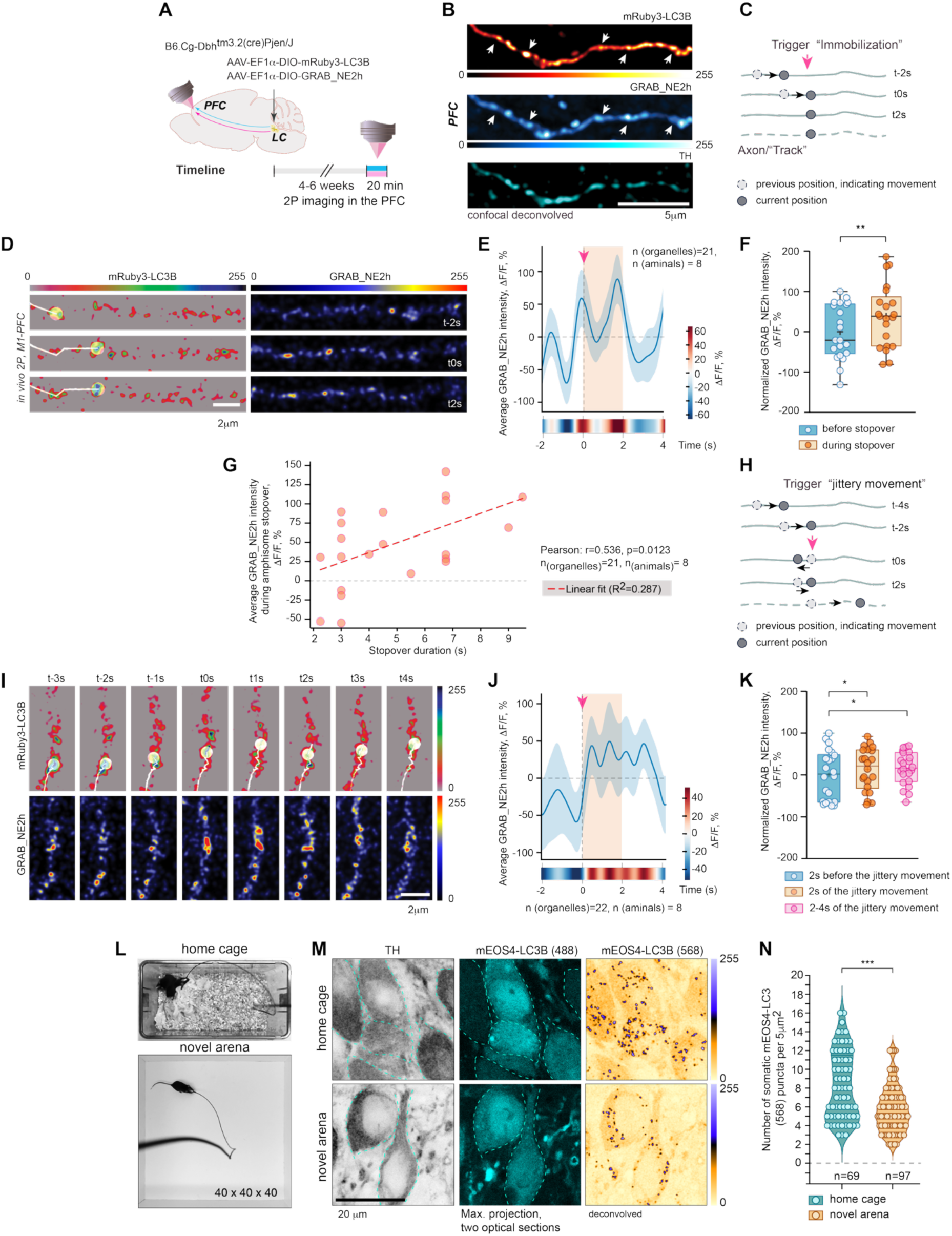
Enhanced *NE* release coincides with localized stopovers and constrained “jittery” amphisome motility in distal *LC* axons *in vivo*. **(A).** AAV-hSyn-DIO-GRAB_NE2h and AAV-EF1α-DIO-mRuby3-LC3B were injected into the *LC*, followed by transcranial window implantation over the PFC/M1. **(B).** Representative confocal images of TH-positive *LC* axons projecting to the PFC showing coexpressing GRAB_NE2h and mRuby3-LC3B. Arrowheads denote regions of increased fluorescence intensity corresponding to LC3B-enriched puncta. **(C).** Schematic illustrating the “trigger”-based selection strategy used to identify unidirectional amphisome immobilization (local stopovers) for automated quantification of NE signal intensity within axonal segments imaged by 2-photon microscopy through a cranial window. **(D).** Representative two-photon imaging frames showing immobilization of mRuby3-LC3B-positive puncta and the corresponding changes in relative GRAB_NE2h fluorescence intensity within distal *LC* axons in the *PFC*, visualized using a pseudocolor lookup table. **(E).** Averaged GRAB_NE2h fluorescence signal (ΔF/F₀%, also indicated by the lookup table) aligned to the “immobilization” as a “trigger”, revealing elevated norepinephrine levels within a 2-s interval surrounding amphisome stopovers. **(F).** Quantification of GRAB_NE2h fluorescence intensity (ΔF/F₀%) 2 seconds before and after amphisome immobilization. P=0.0094 (**\*\***p <0.01), paired *t* test. **(G).** Prolonged amphisome immobilization is associated with increased norepinephrine release, as measured by enhanced GRAB_NE2h fluorescence intensity. Shown is the correlation analysis between amphisome stopover duration and GRAB_NE2h signal intensity. **(H).** Schematic illustrating the “trigger”-based selection strategy used to identify “local jittery movements” for automated quantification of NE signal intensity within axonal segments. **(I).** Representative two-photon imaging frames showing “local jittery movements” of mRuby3-LC3B-positive puncta and the corresponding changes in relative GRAB_NE2h fluorescence intensity within distal *LC* axons in the *PFC/M1*. **(J).** Averaged GRAB_NE2h fluorescence traces (ΔF/F₀%, also indicated by the lookup table) aligned to the onset of localized “jittery” movements used as a “trigger”, revealing elevated NE levels preceding and remaining persistently increased during this distinct motility state *in vivo*. **(K).** Quantification of GRAB_NE2h fluorescence intensity (ΔF/F₀%) measured 2 seconds before and after the “jittery” movements, and 4 seconds during the persistent amphisome “jittery” state. p=0.03 and p=0.0183 respectivly (\**p* < 0.05), paired *t* test. **(L)**. Representative video snapshots acquired during photoconversion experiments under distinct behavioral conditions, including home cage exploration (familiar environment) and exposure to a novel open-field arena. **(M).** Representative confocal images showing photoconverted mEOS4-LC3B-positive puncta (568-nm excitation) and non-photoconverted soluble mEOS4-LC3B fluorescence (488-nm excitation) within LC somata. **(N).** Quantification of somatic photoconverted mEOS4-LC3B-positive puncta across behavioral conditions. Data represent the mean number of somatic photoconverted (“red”) puncta per neuron obtained from six animals per condition: home cage, 69 neurons from 6 animals; novel arena, 97 neurons from 6 animals. \*\**P* < 0.001, unpaired *t* test.

Physiological stimulation of the LC, such as exposure to a novel environment and exploratory behaviour, is associated with increased *LC* firing rates and local NE release (Vankov et al., 1995; Hervé-Minvielle and Sara, 1995; Breton-Provencher et al., 2022). We therefore asked whether activity-dependent confinement of amphisome motility during novelty behavioral context affects their delivery to proximal axonal regions and the soma *in vivo*. To address this question, we performed local fiber-mediated laser photoconversion of mEOS-tagged LC3B within *LC* axons in the *PFC/M1*. Quantitative analysis revealed significantly higher numbers of somatic “red” puncta localized in TH-labeled neurons of the *LC* in animals subjected to photoconversion in their home cages compared with animals exposed to a novel open-field arena during photoconversion (**Fig. 3L-N and S3D-E**). Collectively, our findings support a local signaling mechanism in which heightened NE release during active behavioral states constrains amphisome motility within *LC* axons, limiting retrograde cargo delivery to somatic degradative compartments.

### Activated β2-adrenergic receptors constrain axonal amphisome trafficking through association with SIPA1L2 *in vivo*

Both β1- and β2-adrenergic receptors localize to presynaptic compartments (Langer, 1980; Levin, 1982; Gereau and Conn, 1994; Mizukami, 2004), where they engage Gs-dependent signaling to elevate local cAMP levels and activate PKA, a central regulator of synaptic vesicle dynamics and neurotransmitter release (Nicoll et al., 1994; Weisskopf et al., 1994; Byrne and Kandel, 1996; Gereau and Conn, 1994). We therefore hypothesized that local autoreceptor β-adrenergic signaling may contribute to the confinement of amphisome trafficking observed in *LC* axons under conditions of elevated norepinephrine release. To address this possibility, we first used sparse *LC*-specific GFP labeling combined with three-dimensional axonal reconstruction to visualize distal noradrenergic projections in the *PFC/M1* (**Fig. 1B** and **S1A-B**). Immunohistochemical analysis with antibodies against an extracellular epitope of β2-AR revealed prominent receptor localization in these distal *LC* axons (**Fig. 4A-C; S4A**), supporting a potential presynaptic role for β2-adrenergic signaling in these compartments. The β2-AR contains a conserved C-terminal PDZ-binding motif implicated in receptor trafficking and endosomal sorting through interactions with PDZ-domain scaffold proteins (Hall et al., 1998; Cao et al., 1999). SIPA1L2, a multidomain scaffolding protein associated with LC3B-positive amphisomes (Spilker et al., 2008; Andres-Alonso et al., 2019), contains a class I PDZ domain capable of interacting with C-terminal PDZ-binding motifs (**Fig. S4B**). In *LC* axons, presynaptic β2-ARs were found in close proximity to SIPA1L2, supporting a potential functional interaction *in situ* (**Fig. 4D**). Complementary biochemical and heterologous cell-based assays further confirmed an activation-dependent interaction between β2-AR and SIPA1L2 *in vitro*. Specifically, co-immunoprecipitation assays revealed a robust interaction between SIPA1L2 and β2-ARs, whereas deletion of the receptor C terminus (Δ48) markedly reduced association, indicating a PDZ-binding motif-dependent interaction (**Fig. S4B-D**). Consistent with these biochemical findings, coexpression of FLAG-tagged β2-ARs and mCherry-SIPA1L2 in COS7 cells revealed agonist-dependent co-clustering following stimulation with isoproterenol (**Fig. S4E**). Similarly, surface labeling of FLAG-tagged β2-ARs in primary hippocampal neurons demonstrated spatial association with LC3B-positive amphisomes upon β-ARs agonist isoproterenol treatment (**Fig. S4F-G**). Isoproterenol additionally enhanced colocalization between β2-ARs and SIPA1L2 in cultured hippocampal neurons (**Fig. S4H-I**), suggesting that receptor activation promotes the confinement of SIPA1L2-associated amphisomes at sites of β2-adrenergic signaling. To determine whether activated β2-AR associates with SIPA1L2-positive amphisomes, we employed Nb80, a conformational biosensor that selectively recognizes the active state of the β2-receptor (Rasmussen et al., 2011; Staus et al., 2016). Upon isoproterenol treatment, Nb80, SIPA1L2, and β2-ARs accumulated in proximity at the plasma membrane in COS7 cells (**Fig. S4J**). Moreover, SIPA1L2 co-immunoprecipitated Nb80-GFP from HEK293T lysates expressing β2-AR, supporting an association between SIPA1L2-positive amphisomes and activated β2-AR (**Fig. 4E-F**). Finally, live imaging of SNAP-SiR-LC3B-positive amphisomes in axons expressing FLAG-β2-AR revealed pronounced motility confinement following agonist application, characterized by prolonged immobilization (**Fig. 4G**). Collectively, these findings indicate the existence of a local β2-adrenergic signaling mechanism that constrains axonal amphisome trafficking through PDZ-dependent association with SIPA1L2-positive organelles. Given the close spatial proximity of β2-AR and SIPA1L2 in *LC* axons *in vivo*, it is plausible that the activation-dependent interaction observed in heterologous systems also operates in long-range projecting *LC* axons, although its precise regulatory mechanisms *in vivo* remain to be fully resolved.

**Figure 4.**
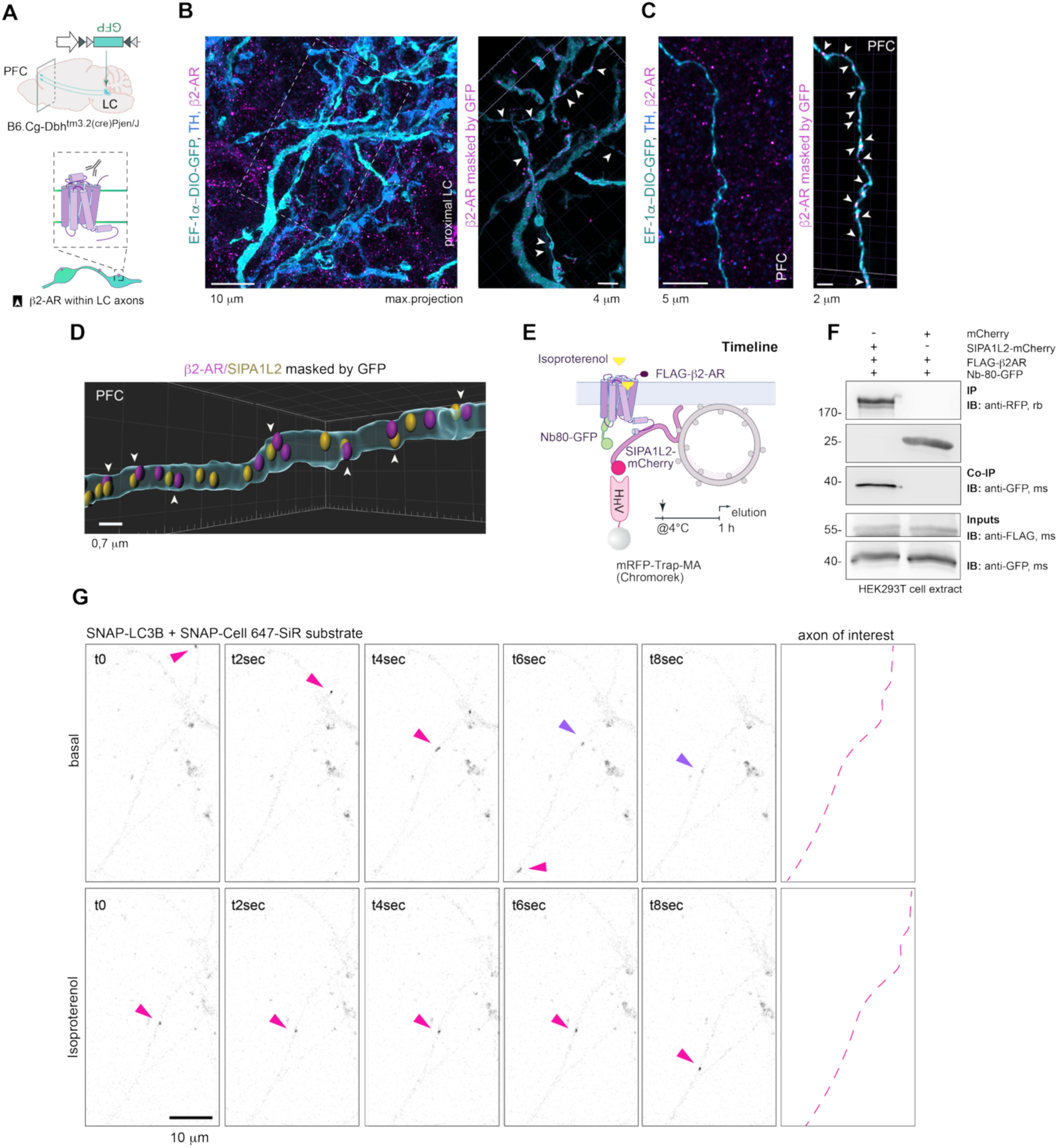
Presynaptic β2-adrenergic receptors constrain amphisome trafficking through the association of activated receptors with SIPA1L2. **(A-C)**. Projection-specific cellular filling followed by three-dimensional masking revealed presynaptic expression of β2-AR within proximal and distal *LC* axons projecting to the prefrontal cortex. **(B).** Representative confocal merged image of GFP-filled *LC* processes (cyan), positive for tyrosine hydroxylase (TH; blue). Magnified regions of interest show β2-AR puncta within GFP-positive axons (arrowheads). **(C).** Representative confocal merged image of a GFP-labeled, TH-positive *LC* axon within the prefrontal cortex. Magenta indicates β2-AR immunoreactivity. Arrowheads denote receptor-positive puncta localized within GFP-filled axonal segments. **(D).** SIPA1L2-positive puncta are frequently detected in proximity to presynaptic β2-adrenergic receptors within GFP-labeled *LC* axons projecting to the PFC (arrowheads). **(E, F).** SIPA1L2 associates with activated β2-AR as detected using the conformational nanobody biosensor Nb80. **(E).** Schematic representation of the experimental workflow. **(F).** Immunoblot analysis showing association of SIPA1L2, but not mCherry alone, with activated β2-AR identified by Nb80 binding. Cells were treated with isoproterenol for 30 min before lysis. **(G).** Representative confocal time-lapse imaging of an axon from a primary neuron co-expressing FLAG-tagged β2-AR and SNAP-LC3B, before and after isoproterenol treatment. Images were smoothed and denoised using the despeckle function in Fiji (ImageJ). Dashed line indicates the axon of interest.

### Reduced NE signaling and Gi-mediated inhibition enhance directional amphisome trafficking in *LC* axons

Having established that amphisome confinement is associated with elevated local NE signaling, we next investigated whether reductions in local NE promote directional amphisome transport. Given the pronounced heterogeneity of amphisome trafficking states (**Fig. 1**), we hypothesized that transitions between motility states are dynamically coupled to fluctuations in local axonal NE signaling. Specifically, we examined whether relative reductions in local NE facilitate amphisome mobilization by a local regulatory mechanism. Consistent with this hypothesis, spatiotemporal tracking analysis of amphisomes in *LC* axons *in vivo* revealed that episodes of sustained unidirectional movement were associated with marked local decreases in GRAB_NE2h fluorescence compared with the previous immobile states, indicating that reduced local NE signaling favors unidirectional amphisome mobilization (**Fig. 5A-D; 3A-B; Fig. S3A-C**). Presynaptic α2A-adrenergic autoreceptors suppress NE release via Gi-coupled inhibition of the cAMP/PKA pathway (Park et al., 2009; Harris et al., 2018; Jewell-Motz et al., 1998; Brown et al., 2022) and are thought to contribute to autoregulatory modulation of *LC* activity during low-arousal states, including sleep- associated neuromodulatory conditions (Silverman et al., 2025). We therefore hypothesized that Gi signaling may promote directional amphisome trafficking by reducing PKA-dependent immobilization and facilitating retrograde transport toward somatic degradative compartments (**Fig. 5E**). To test this possibility, we combined *in vivo* two-photon imaging of Cre-inducible LC3B-labeled amphisomes through a cranial window (**Fig. 1D-E**) with chemogenetic suppression of *LC* neuronal activity using the Gi-coupled DREADD receptor hM4D(Gi) (AAV-hSyn-DIO-hM4D(Gi)-mCherry), a well-established approach for mimicking inhibitory α2A-adrenergic receptor signaling (Perez et al., 2020; McCall et al., 2015; Wagatsuma et al., 2018; **Fig. 5A and S5A**). Systemic administration of the selective DREADD agonist JHU37160 (J60; Bonaventura et al., 2019) significantly increased the average speed of amphisomes within distal *LC* axons relative to baseline conditions, as quantified in the same axonal segments before and after treatment, enabling a direct within-axon comparison (**Fig. 5F-H**). We next examined whether Gi activation unifies amphisome directionality by suppressing PKA-dependent organelle immobilization. Under baseline conditions, mobile amphisomes displayed comparable proportions of unidirectional and jittery trajectories (**Fig. 5I-J**). Strikingly, Gi activation reduced the fraction of jittery amphisomes to ∼34% and increased the average velocity by ∼50%, producing a pronounced shift toward highly directional transport *in vivo* (**Fig. 5J-K**). Consistent with previous observations that LC3B-positive amphisomes segregate into distinct trafficking populations under steady-state *LC* activity (**Fig. 1I**), PCA revealed a marked reorganization of transport dynamics following J60 treatment (**Fig. 5L-M**). At baseline, amphisomes exhibited slow and fast jittery trajectories and slow unidirectional trajectories, whereas Gi activation promoted predominantly unidirectional transport across slow, fast, and extra-fast velocity classes (**Fig. 5M**). Collectively, these findings demonstrate that Gi signaling, likely through suppression of cAMP/PKA activity, reduces amphisome immobilization and non-directional jittery switching, thereby promoting rapid and highly directional trafficking along distal *LC* axons. Together, these data indicate that both reduced local norepinephrine signaling and Gi-mediated inhibition converge to mobilize axonal amphisomes, increase averaged transport velocity, and reinforce directional retrograde trafficking *in vivo*.

**Figure 5.**
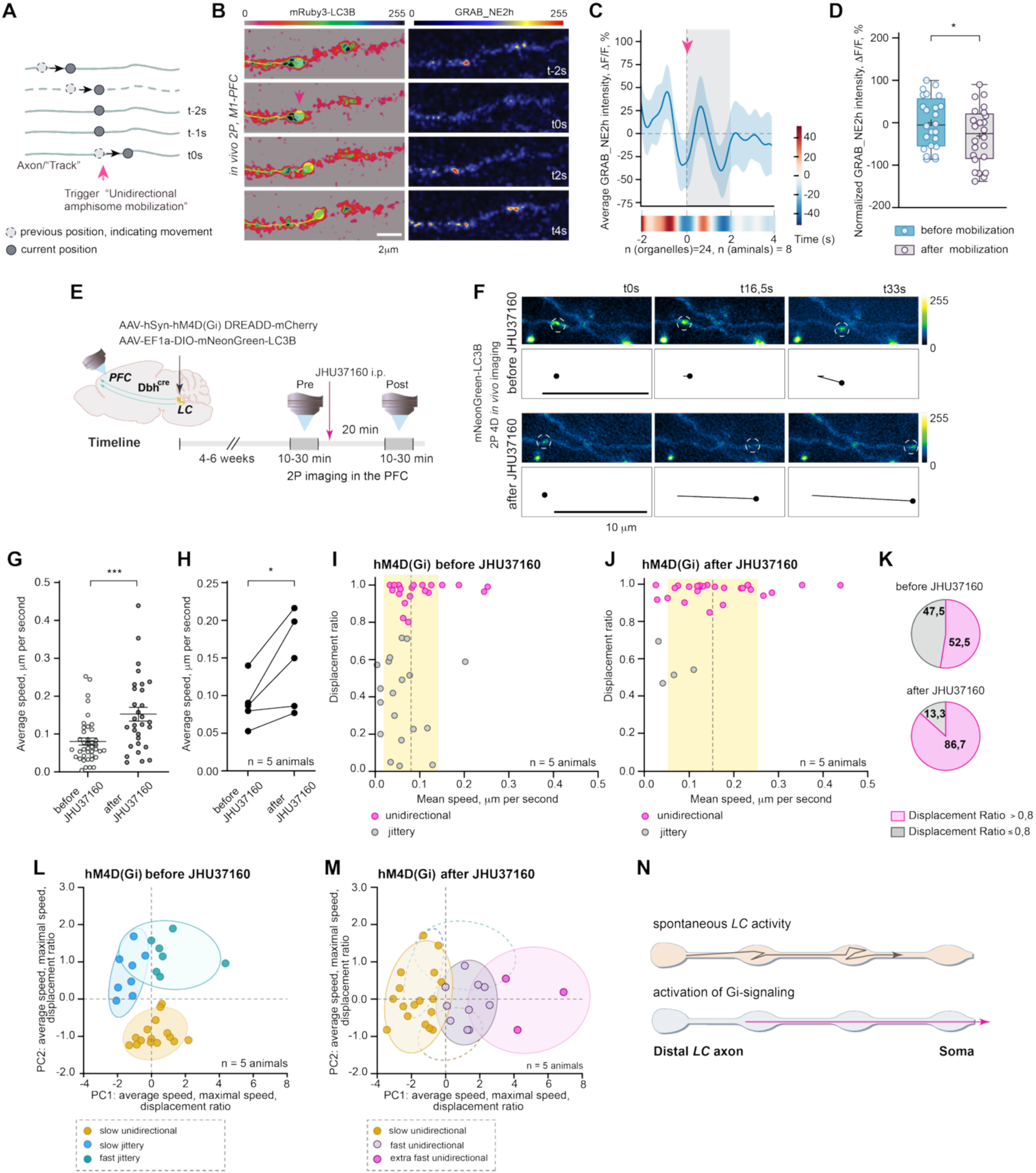
Reduced norepinephrine level and Gi-mediated inhibition enhance processive amphisome trafficking in *LC* axons. **(A)**. Schematic representation of trigger-based selection of amphisome mobilization events for quantification of local NE dynamics in *LC* axons *in vivo*. **(B-D)**. Representative two-photon imaging sequences **(B)** illustrating mobilization of mRuby3-LC3B-positive puncta together with a concomitant reduction in GRAB_NE2h fluorescence within distal *LC* axons, displayed using a pseudocolor lookup table **(C)**. **(D).** Normalized mean GRAB_NE2h fluorescence (ΔF/F₀%) aligned to amphisome mobilization (“trigger”), showing a significant decrease in NE levels during and after mobilization. p=0.0422 (**\***p < 0.05), unpaired *t-test* **(E).** Schematic illustrating the experimental timeline for *in vivo* assessment of amphisome trafficking in *LC* axons before and after chemogenetic activation of Gi-coupled signaling. **(F)**. Representative *in vivo* two-photon time-lapse frames acquired before and after intraperitoneal (ip) administration of JHU37160. Dashed circles indicate automatically detected and TrackMate-tracked mNeonGreen-LC3B-positive amphisomes (ImageJ), with manual annotation. The schematic depicts the displacement of a representative amphisome before and after JHU37160 treatment. **(G)**. Average amphisome speed across *LC* axons before and after Gi activation. Data are mean ± SEM; p=0.001 (***), unpaired *t-test*. n, total number of mobile amphisomes from 5 animals. **(H).** Quantification of amphisome speed averaged across multiple organelles within the same *LC* axons before and after JHU37160 administration. p=0.0381 (*p < 0.05), paired *t-test*. n=5 animals. **(I, J).** Cluster analysis of individual amphisome trajectories before **(I)** and after **(J)** JHU37160 treatment. Each point represents a single trajectory; the grey dashed line indicates mean trafficking velocity, and the yellow-shaded area denotes the standard deviation. Gi activation broadens the distribution of mean trafficking speeds **(J)**. Magenta circles represent unidirectional amphisomes (displacement ratio ≥ 0.8); grey circles represent jittery vesicles. **(K).** Pie chart quantifying the proportion of amphisomes exhibiting unidirectional vs. jittery motility before and after JHU37160 treatment. Gi activation decreased the proportion of jittery amphisomes by ∼34% and increased unidirectional trafficking (n = number of amphisomes analysed before and after treatment, from 5 animals per group). **(L, M)**. PCA-based visualization of amphisome trafficking states before and after JHU37160 administration. **(L)**. Three clusters were identified: slow unidirectional (yellow), slow jittery (blue), and fast jittery (cyan). Pre-treatment trajectories were combined with those from spontaneous LC activity (Fig. 1I) to generate a unified baseline distribution. **(M)**. Following chemogenetic Gi activation, clusters shift to slow unidirectional, fast unidirectional, and extra-fast unidirectional states. Ellipses indicate cluster distributions; dashed lines denote PCA axes representing feature variance. Post-treatment clusters are overlaid on baseline distributions for comparison. **(N)**. Schematic illustrating Gi signaling–dependent reduction of amphisome immobilization and jittery states, promoting unidirectional trafficking and increased transport velocity in distal *LC* axons.

## DISCUSSION

Neuronal autophagy has been studied extensively, but how autophagy-related processes in specialized axons are regulated *in vivo,* and whether they are modulated by neuromodulatory transmission, remains largely unknown. Disruption of retrograde trafficking and maturation of autophagy-related organelles, including amphisomes, can lead to their pathological accumulation within dystrophic axonal swellings, a hallmark of multiple neurodegenerative disorders (Lie et al., 2021). Because amphisomes form locally within distal axons (Andres-Alonso et al., 2026), whereas fully mature lysosomes remain largely restricted from distal neuronal processes in the mature brain (Lie et al., 2021), efficient retrograde transport toward somatic degradative compartments in *LC* neurons is likely to require precise local mechanisms coordinating cargo delivery, potentially governed by synaptic activity and behavioral state. In the present study, we demonstrate that SIPA1L2-positive amphisomes are abundant within long-range projecting *LC* axons and are frequently localized to axonal varicosities resembling presynaptic boutons (**Fig. 1 and S1**). To directly test whether distally generated amphisomes undergo long-range retrograde transport, we employed fiber-mediated photoconversion and detected photoconverted amphisomes either in proximal neuronal processes and within the somata themselves (**Fig. 2**).

Two-photon time-lapse imaging through a cranial window in awake mice, combined with principal component analysis of trafficking dynamics, identified three major motility states of axonal amphisomes: slow jittery, fast jittery, and slow unidirectional populations. Notably, most mobile amphisomes exhibited transient immobilization events lasting several seconds, thereby slowing progressive retrograde cargo delivery. Previous work demonstrated that stationary pauses of SIPA1L2/LC3-positive amphisomes are regulated by cAMP/PKA-dependent mechanisms controlling both dynein motor engagement and organelle processivity (Andres-Alonso et al., 2019). These mechanisms include modulation of amphisome interactions with the dynein motor complex through the adaptor protein snapin (Andres-Alonso et al., 2019; Cai et al., 2010; Xie et al., 2015; Cheng et al., 2015), as well as regulation of motor processivity through the GAP activity of SIPA1L family proteins associated with the outer membrane of LC3-positive organelles (Andres-Alonso et al., 2019; Goldsmith et al., 2022).

### Regulation of amphisome trafficking *in vivo*

Building on these observations, we further hypothesized that prolonged local presynaptic norepinephrine signaling associated with elevated auto-cAMP/PKA activity during distinct behavioral states promotes confinement of amphisome motility and restricts retrograde transport, whereas suppression of local cAMP/PKA signaling, often associated with low *LC* activity, facilitates directional trafficking and thereby enhances somatic cargo delivery.

We next monitored local NE dynamics in *LC* axons *in vivo* using the genetically encoded fluorescent sensor GRAB_NE2h, which enables rapid and selective reporting of NE fluctuations (Feng et al., 2024). Intriguingly, comparison of NE sensor dynamics with amphisome motility revealed a clear temporal coordination between both processes. Motility states that restrict processive transport, including transient immobilization and localized “jittery” dynamics, likely reflecting PKA-dependent modulation of dynein processivity and intermittent motor engagement, were consistently associated with elevated local NE signaling *in vivo* (**Fig. 3**). Prolonged stationary phases of amphisomes were associated with sustained NE elevations, suggesting a local feedback relationship between organelle dynamics and NE signaling (**Fig. 3G**). Beyond their degradative role, amphisomes have been proposed to function as signaling platforms that retain signaling competence during retrograde transport prior to lysosomal fusion (Kononenko et al., 2017; Andres-Alonso et al., 2019; Karpova et al., 2025; Andres-Alonso et al., 2026), raising the possibility that immobilized amphisomes contribute to local modulation of presynaptic function in *LC* axons. In rodents, exposure to novel environments robustly activates *LC* neurons and transiently elevates forebrain norepinephrine levels, providing a naturalistic, behaviorally defined condition of high presynaptic NE signaling against which trafficking dynamics can be assessed in the intact, awake animal (Vankov et al., 1995; Liu et al., 2015; Takeuchi et al., 2016; Wagatsuma et al., 2018). Consistent with this idea, higher *LC* activity during anxiety-like behavior was associated with reduced numbers of amphisomes reaching somatic compartments following photoconversion in *PFC/M1*-projecting axons (**Fig. 3L-N**).

What might be the advantage of integrating degradative and signaling functions within a single organelle in *LC* axons? In the present study, combining projection-specific labeling approaches, we show that LAMP2A-positive lysosomal structures are exceedingly sparse within distal *LC* axons (**Fig. 2**). We speculate that such an organization enables tight coordination of proteostatic and synaptic demands at sites of elevated neuronal activity. Given that NE signaling operates predominantly through volume transmission from multiple axonal varicosities rather than discrete synapses, this integrative arrangement may provide distributed local feedback control across functionally engaged release domains. In this context, signaling-competent amphisomes may function not only as carriers of degradative cargo, but also as local sensors and regulators of presynaptic activity, dynamically coupling neuronal activity states to axonal proteostasis and cargo trafficking.

How can this be achieved at the molecular level? An important unresolved question concerns the signaling capacity of *LC* amphisomes. In previous work, we identified BDNF/TrkB-positive signaling amphisomes in hippocampal neurons (Andres-Alonso et al., 2019; Karpova et al., 2025), and BDNF/TrkB signaling has likewise been reported in *LC* neurons (Matsunaga et al., 2004; Traver et al., 2006; Liu et al., 2015; Suto et al., 2019). Amphisome immobilization requires cAMP/PKA signaling and dissociation of the dynein adaptor snapin from the motor complex (Zhou et al., 2012; Cheng et al., 2015; Di Giovanni and Sheng, 2015; Andres-Alonso et al., 2019). Here, we show that β2-adrenergic receptors, whose activation engages cAMP/PKA signaling, are expressed presynaptically in distal *LC* axons. Moreover, β2-AR and SIPA1L2 were frequently detected in close proximity within these axons, and SIPA1L2 preferentially associated with activated β2-adrenergic receptors identified using a nanobody-based conformational biosensor (Rasmussen et al., 2011; Staus et al., 2016). In this framework, activated β-AR, established regulators of presynaptic vesicle dynamics and neurotransmitter release (Nicoll et al., 1994; Weisskopf et al., 1994; Byrne and Kandel, 1996; Gereau and Conn, 1994; Chen et al., 2011), may restrict processive retrograde transport by docking SIPA1L2-positive amphisomes near enhanced sites of NE release, thereby reinforcing local neurotransmission and presynaptic signaling. At the same time, the multidomain scaffolding properties of SIPA1L2 raise the possibility that *LC* neurons harbor multiple signaling amphisome populations associated with distinct receptor and signaling complexes beyond TrkB-containing organelles. In this context, it is tempting to speculate that heightened release activity and elevated metabolic demand, which are expected to increase protein damage, may necessitate enhanced formation and turnover of amphisomes to ensure efficient proteostatic clearance.

### *LC* inhibition promotes amphisome mobilization, and sleep-associated neuromodulatory states accelerate and unify directional trafficking in *LC* axons

Elevated NE signaling through presynaptic β-adrenergic receptors slows amphisome trafficking by promoting prolonged distal immobilization or localized “jittery” motility states. In contrast, amphisome mobilization is initiated during phases of reduced local NE release (**Fig. 5 and Fig. 6**). Along these lines, selective activation of Gi-coupled α2-adrenergic autoreceptors, known to define low-arousal *LC* states and mimic REM phase in a sleep-associated neuromodulatory conditions (Hansen & Manahan-Vaughan, 2015; Nguyen & Connor, 2019; Lim et al., 2010; Lemon et al., 2009; Maity et al., 2015; Silverman et al., 2025), shifted amphisome trafficking toward a more processive state. Chemogenetic Gi activation via hM4D(Gi) and JHU37160 not only increased the fraction of unidirectional amphisomes (52.5% to 86.7%) but also reorganized trafficking states, replacing slow and fast “jittery” populations with predominantly unidirectional modes and increasing overall transport velocity by >50%. In summary, we propose a model in which behavioral state and neuromodulatory tone bidirectionally regulate amphisome dynamics in *LC* axons and shape axonal-to-somatic amphisome delivery. During wakefulness, in axons with elevated NE signaling, transient immobilization of amphisomes at axonal varicosities coordinates local presynaptic function and activity-dependent cargo handling (**Fig. 6**). In contrast, during sleep-associated low-NE states, reduced cAMP/PKA signaling favors uninterrupted retrograde transport and efficient delivery of autophagic cargo to somatic degradative compartments.

**Figure 6.**
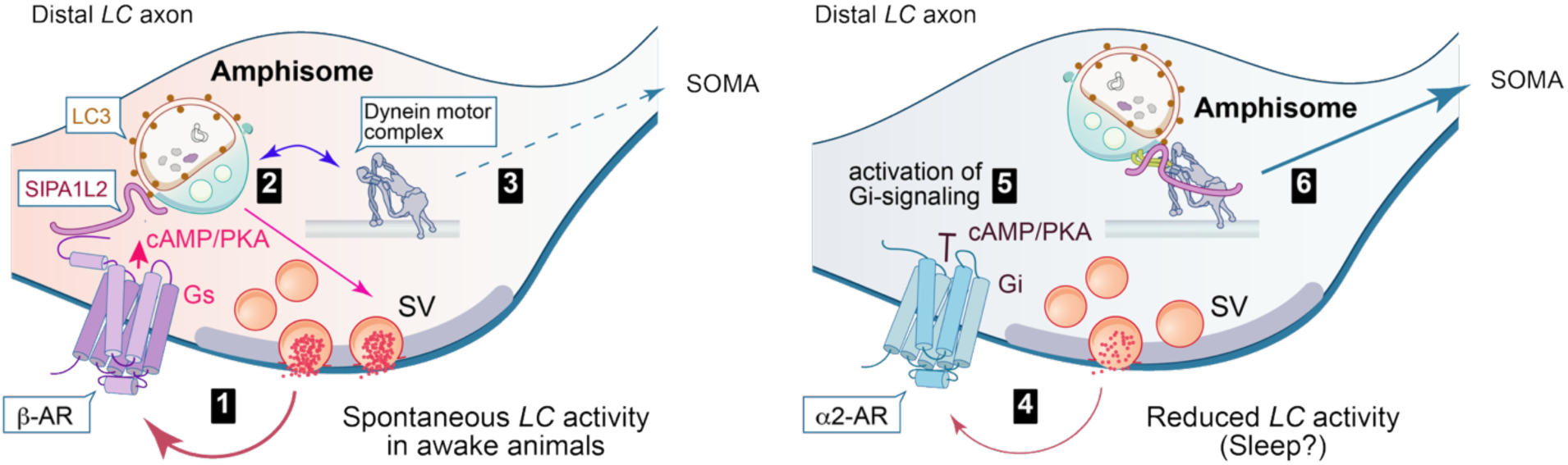
Model of activity-dependent regulation of axonal amphisome trafficking in which behavioral state and neuromodulatory tone bidirectionally control amphisome dynamics in *LC* axons and shape axonal-to-somatic amphisome delivery. Behavioral states associated with elevated local axonal NE release (1) activate presynaptic β-adrenergic autoreceptors, leading to engagement of cAMP/PKA signaling and the induction of amphisome confinement states (2), including immobilization and jittery, non-processive dynamics, possibly via association of b2AR with SIPA1L2. These motility states limit efficient axonal-to-somatic delivery of autophagic cargo (3). In contrast, during sleep-associated low-NE states (4), reduced cAMP/PKA signaling (5) favors uninterrupted retrograde transport of amphisomes along *LC* axons, enabling efficient delivery of autophagic cargo to somatic degradative compartments (6).

### Perspectives

Our study establishes a conceptual framework linking neuromodulatory state, axonal autophagy, and neuronal proteostasis in vivo. The brainstem *LC*, whose neurons provide widespread projections to cortical, hippocampal, thalamic, and subcortical regions (Swanson and Hartman, 1975; Schwarz and Luo, 2015), represents a key model in which long-range axons must coordinate neuromodulatory signaling with continuous intracellular cargo trafficking. Given that *LC* degeneration is among the earliest events in several neurodegenerative disorders, impaired activity-dependent regulation of amphisome trafficking may contribute to defective proteostasis well before overt neuronal loss. Future studies should determine whether disruption of this neuromodulatory transport mechanism represents a convergent pathway underlying selective vulnerability of long-range projecting neurons in neurodegenerative disease.

## MATERIALS AND METHODS

### EXPERIMENTAL MODEL

#### Animals

All animal procedures were conducted in accordance with the Directive of the European Parliament and Council on the protection of animals used for scientific purposes (2010/63/EU). Experiments requiring authorization (§7 VersTierV, German Animal Welfare Laboratory Animal Ordinance) were approved by the competent authority of the federal state of Saxony-Anhalt, Germany (Landesverwaltungsamt für Verbraucherschutz und Veterinärangelegenheiten, Referat 203; State Office for Consumer Protection and Veterinary Affairs, Department 203; license number 42502-2-1638 LIN) and performed in compliance with these regulations. B6.Cg-Dbh^tm3.2(cre)Pjen/J mouse line (Strain #:033951 RRID: IMSR_JAX:033951) was obtained from the Jackson Laboratory and maintained at the animal facility at the Leibniz Institute for Neurobiology (LIN), Magdeburg, Germany. Animals were housed under institutional animal welfare guidelines, and all surgical procedures were performed at 10-12 weeks of age.

## METHOD DETAILS

### Cloning of viral constructs and viral partial production

Open reading frames (ORFs) for mNeonGreen-LC3B, mRuby3-LC3B, and mEOS4-LC3B were synthesized as gBlocks (Integrated DNA Technologies) and subcloned into the AAV-EF1α-DIO backbone (Addgene #50462) using SgsI/AscI-NheI restriction sites. AAV particles expressing fluorescently tagged LC3B were produced as DJ and AAV2/9 serotypes. Additional AAV particles were obtained as ready-to-use preparations from Addgene or the Zurich Viral Vector Facility (see key resource table for details). For AAV production, the standard iodixanol gradient ultracentrifugation protocol provided by Addgene was used with minor modifications. Briefly, HEK293T cells were plated in three 150-mm petri dishes and co-transfected the following day with the expression plasmid, the packaging plasmid (pAAV-DJ or pAAV2/9), and pHelper at a 1:1:1 molar ratio using polyethylenimine (PEI). Cells were harvested three days post-transfection, washed with 10 mL prewarmed PBS, resuspended in 10 mL PBS, pooled from the three dishes, and centrifuged at 1,000 × g for 30 min at 4 °C.

The cell pellet was stored at -70°C overnight, while the supernatant was transferred into fresh tubes. A PEG precipitation was performed by adding 4 g of polyethylene glycol and 2,32 g of NaCl per 40 ml of supernatant, followed by shaking for 3 h in an overhead shaker at 4 °C and then resting overnight at 4 °C. The solution was subsequently centrifuged at 3,000 × g for 30 min at 4 °C. The resulting pellet was resuspended in 2 ml of Tris-HCl buffer (50 mM TRIS base, 150 mM NaCl, pH 8.5). The initial cell pellet was resuspended in 3 mL of the same TRIS-HCl buffer and subjected to three freeze-thaw cycles. The lysate was combined with the PEG-precipitated solution and treated with Benzonase (50 U/mL) for 1 h at 37 °C to degrade residual nucleic acids. The mixture was clarified by centrifugation at 8,000 × g for 30 min at 4 °C, and the supernatant was filtered through a 0.2 μm membrane before AAV purification. AAV particles were purified by ultracentrifugation in an iodixanol step gradient consisting of 8 ml 15% (prepared in 1 M NaCl/PBS-MK buffer: 5.84 g NaCl, 26.3 mg MgCl₂, and 14.91 mg KCl in 100 ml PBS), 6 ml 25%, 5 mL 40%, and 5 mL 60% (both prepared in PBS-MK buffer: 26.3 mg MgCl₂ and 14.91 mg KCl in 100 ml PBS). The 25% and 60% fractions were supplemented with phenol red for visualization. Gradients were centrifuged in 38.6 mL Ultra-Clear tubes (SW32 Ti rotor, Beckman) at 170,000 × g for 16h at 4°C. For washing and concentration of AAV particles, the 40% fraction was applied to centrifugal filter units preconditioned with 0.1% Pluronic F-68 in PBS for 10 min at room temperature. Filters were subsequently rinsed with 0.01% and 0.001% Pluronic F-68 in PBS containing 200 mM NaCl, followed by iterative centrifugation (3,000–3,500 rpm, 4 °C) until the viral preparation was concentrated to 200-250 μL. Viral concentrates were recovered by rinsing the filter walls and stored at 4 °C for short-term use (≤2 weeks) or aliquoted and stored at –80 °C for long-term use. Viral titres were determined by quantitative PCR using the following primers: forward ITR primer, 5′-GGAACCCCTAGTGATGGAGTT-3′ and reverse ITR primer, 5′-CGGCCTCAGTGAGCGA-3′ (Aurnhammer et al., 2012).

### AAV injection into the *Locus coeruleus*

For stereotactic injections, animals were anesthetized with ketamine/xylazine (100/5 mg/kg, intraperitoneally) and placed in a stereotactic frame. The skull was exposed and cleaned, and the bregma and lambda landmarks were identified. For targeting the locus coeruleus, stereotaxic coordinates relative to bregma were anterior-posterior (AP) -5.4 mm, medial–lateral (ML) ±0.93 mm, and dorsal-ventral (DV) -3.6 mm (Allen Mouse Brain Atlas; available at http://mouse.brain-map.org). A small craniotomy (0.1– 0.2 mm) was drilled, and a Hamilton syringe mounted on a nanoliter (nL) injector was lowered slowly (≤0.2 mm/s). After a 5-minute pause, 500 nL of virus was injected at 75 nL/min (Cearley and Wolfe, 2007). The needle remained in place for 10 min post-injection before being withdrawn slowly (0.1-0.2 mm/s) to minimize tissue damage. The incision was closed with silk sutures and reinforced with dental cement. Animals were monitored during recovery and received postoperative analgesia (carprofen, 5 mg/kg, s.c.). Surgical sites were inspected daily, with wound healing confirmed 3 days post-surgery, and animals were monitored until tissue collection.

### Optical fiber implantation and optical fiber-mediated photoconversion in the home cage versus the open arena

Following viral expression, an optical fiber was implanted above the *mPFC* to enable in vivo photoconversion. Anesthesia and preparatory procedures were identical to AAV injections. A 300-µm craniotomy was drilled above the right *mPFC* (AP +1.8 mm, ML +0.3 mm relative to bregma), and a 300-µm fiber cannula mounted in a ceramic ferrule was lowered to DV−2.2 mm at 1 μm/min and secured with dental cement mixture. Wounds were sutured, and animals received identical postoperative care and analgesia. Four to six weeks post-surgery, the implanted fiber was coupled to a 405 nm laser with 5 mW output through a flexible fiber cord (300 μm core, ∼85% transmission efficiency), allowing unrestricted movement of the animal. Photoconversion of mEOS-LC3B in the *mPFC* was performed in awake mice under two conditions: a familiar environment (home cage) and a novel environment (40 × 40 × 40 cm white-walled arena; McCall et al., 2017). Identical stimulation parameters were applied across contexts (405 nm, 5 mW, 600 s total, 1 s pulses with 10 s intervals). Mice were video recorded throughout the procedure to monitor locomotion and exploration, enabling correlation of behavioral state with photoconversion outcomes. Six hours after photoconversion, mice were anesthetized and perfused with 4% PFA. Brains were dissected, postfixed, and sectioned (35-40 μm) under low-light conditions to minimize stochastic photoconversion. Coronal *LC* sections were collected in PBS and processed for immunohistochemistry. Confocal microscopy was used to quantify photoconverted LC3B puncta within *LC* somata.

### Transcranial window implantation

For cranial window implantation, animals were prepared as described above. A larger portion of the skull was exposed, and connective tissue was gently removed to prepare the bone surface. A 5 mm circular craniotomy was drilled over the prefrontal-motor cortex region with continuous saline irrigation to prevent heating (Zuluaga-Ramirez et al., 2015; Goldey et al., 2014). The skull flap was removed carefully without disrupting the dura, and a 5 mm glass coverslip was placed over the exposed cortex. The coverslip was sealed with cyanoacrylate adhesive and stabilized with dental cement (Xu et al., 2007; Mostany & Portera-Cailliau, 2008; Goldey et al., 2014; Holtmaat et al., 2009; Yang et al., 2010). A custom-made metal headplate was affixed over the cranial window using SuperBond and dental cement to enable stable head fixation during two-photon imaging. Animals recovered under standard postoperative care.

### Two-photon imaging *in vivo* of *LC* axons projecting to *PFC/M1*

Two to four weeks after AAV transduction and window implantation, *in vivo* two-photon imaging was performed using a Bergamo microscope (Thorlabs, Newton, NJ, USA) equipped with a tunable Ti: sapphire laser (Chameleon Vision S, Coherent Inc.) and a fixed-wavelength laser (HighQ-2, Spectra-Physics Inc.). Both lasers were aligned to the same foci and operated simultaneously to enable dual-colour imaging in 4D (XYZT), whereas the wavelength of the Vision S laser was used in the range between 920 nm and 960 nm for green-emitting fluorophores (mNeonGreen-LC3B, GRAB_NEh), and the HighQ-2 laser (wavelength 1040 nm) was used for red fluorophores (mRuby3-LC3B). Actual power for in-vivo imaging with each laser was set to be less than 30 mW to minimize photobleaching or damage in long imaging sessions. Prior to imaging, animals were handled and habituated to the setup for five days to reduce stress and ensure their comfort during the procedure. On the first day of imaging, animals were briefly anesthetized with isoflurane (30 seconds) and head-fixed under the objective. Imaging areas were first localized with a 4x objective under one-photon excitation (488 nm LED, X-CITE 200) and detected with a camera (1500M-GE, a 1.4 MP) to capture an overview image and adjust the focal plane for the region of interest. High-resolution data were acquired with a 40x water-immersion objective (NA 0.8) in two-photon mode. Distal axons in the *mPFC/M1* region were imaged with a digital zoom of 10-13x. Optical sectioning was performed in 1 µm Z-steps, with 30 Hz frame rate, collecting 100-150 repeated stacks per axon. Fifteen frames were averaged per plane, resulting in acquisition times of 20-30 min with ∼2-3 µm resolution for axon thickness. For GRAB_NE2h imaging, data were acquired in 3D (XYT) at 30 Hz for 20 min to ensure continuous recording of norepinephrine signals. Data and metadata were saved automatically for subsequent analysis.

### Combined two-photon imaging and chemogenetic silencing

For combined two-photon imaging and chemogenetic activation of Gi-signaling in *LC* neurons, B6.Cg-Dbh^tm3.2(cre)Pjen/J mice received stereotactic co-injections of AAV-EF1a-DIO-mNeonGreen-LC3B and AAV-EF1a-DIO-hM4D(Gi)-mCherry (Armbruster et al., 2007) into the *LC*, followed by implantation of a transcranial window and headplate, as described above. Axonal stretches were imaged sequentially under basal conditions and following chemogenetic activation of Gi signalling. The DREADD agonist JHU37160 (Bonaventura et al., 2019) was administered subcutaneously at 0.5 mg/kg immediately after the initial imaging session. After a 20-30 min post-injection interval to allow sufficient blood-brain barrier penetration (Lawson et al., 2024), the same axonal stretch was re-imaged. This design enabled direct comparison of LC3B-positive vesicle trafficking during spontaneous activity versus Gi*-*signalling activation. Two-photon imaging was conducted using the parameters described above. Z-stacks were acquired with a 1 μm step size, 10-13x digital zoom, and 30 Hz acquisition rate, averaging 15 frames per Z-plane to reduce motion artifacts. For each axon, 100-150 repeated stacks were collected.

### Immunohistochemistry

Animals were perfused transcardially with PBS followed by 4% PFA. Extracted brains were post-fixed in 4% PFA for at least 8 hours and cryoprotected in 30% sucrose for 48–72 hours. Coronal brain sections (40 μm) were prepared using a freezing microtome (HM440E, Microm). Heat-based antigen retrieval was performed by incubating sections in 10 μM sodium citrate (pH 9, adjusted with 5M NaOH) for 30 min at 80 °C. Sections were then permeabilized in 0.2% Triton X-100 in PBS for 1 hour and blocked in blocking buffer (2% glycine, 0.2% gelatine, 2% BSA, and 50 mM NH_3_Cl (pH 7.4)). Primary antibodies were incubated for 24-48h at 4°C in blocking buffer, followed by three 10-minute PBS washes and incubation with fluorophore-conjugated secondary antibodies for 1 hour at room temperature. After a final series of PBS washes and a brief rinse in distilled water, sections were mounted with Mowiol 4-88 (Merck Chemicals GmbH) and imaged using confocal microscopy.

### Confocal laser scanning microscopy and image analysis

Confocal images of fixed tissue samples were acquired using a Leica TCS SP8 STED 3X confocal laser scanning microscope equipped with a 405 nm diode laser and a white-light laser (WLL) (Leica Microsystems, Mannheim, Germany). For imaging *LC* neurons and their terminals in the *PFC*/*M1*, 63×/1.4 NA (HCX PL APO, Leica) or 100×/1.4 NA (HC PL APO, Leica) oil-immersion objectives were used. Images were acquired at 400-600 Hz with a lateral resolution of 1024 × 1024 pixels and a Z-stack step size of ∼160-180 nm employing Hybrid (HyD) detectors.

### Image analysis

#### Amphisome detection and characterization analysis in Z-stack acquired *in vivo*

Image processing and analysis were performed using ImageJ/FIJI, following a multi-step workflow optimized for amphisome quantification. Pre-processing included sequential digital filters: rolling ball background subtraction (radius optimized to feature size), median filtering (2-3-pixel radius), and Gaussian smoothing (σ = 1-2 pixels) to reduce noise while preserving biological structures and improving signal-to-noise ratio (Schindelin et al., 2012; Berg et al., 2019). Contrast enhancement employed histogram-based methods, with parameters documented to maintain data integrity (Aaron & Chew, 2021). Regions of Interest (ROIs) were selected based on standardized anatomical criteria, incorporating morphological features and fluorescence intensity profiles. Automated ROI detection algorithms, supplemented by manual verification, ensured consistent selection across experimental conditions. Multi-channel analysis isolated specific vesicle populations via channel splitting and colocalization, with Max-Entropy thresholding applied using machine learning-optimized parameters for accurate segmentation across the full Z-stack (Arganda-Carreras et al., 2017; Wiesmann et al., 2014). Amphisome quantification utilized the StarDist deep learning-based object detection algorithm, trained on manually annotated vesicle datasets. StarDist employs a U-Net architecture with star-convex polygon representation, enabling accurate separation of closely packed vesicles while preserving morphological integrity (Schmidt et al., 2018; Weigert et al., 2020; Völker et al., 2020; Fazeli et al., 2021). The pipeline generates vesicle counts per ROI, size distributions (μm²), and shape parameters, with automated circularity measurements (4π × area/perimeter²) and variance statistics. Quality control included merging thresholded images with pre-processed data for visual verification. This standardized workflow ensures reproducible vesicle detection and characterization with high sensitivity and specificity.

#### Image processing and amphisome tracking analysis

Time-lapse *in vivo* imaging of amphisome dynamics in axons requires precise processing to preserve spatial and temporal accuracy. Motion correction was performed in ImageJ/FIJI using the FAST4DREG plugin, which incorporates the NanoJ-Core algorithm for sub-pixel registration across all dimensions (x, y, z, t) (Laine et al., 2019; Culley et al., 2018). Motion correction parameters were optimized iteratively for each dataset to maximize registration accuracy while minimizing artifacts, enabling reliable tracking of small, rapidly moving vesicles in living tissue. Following motion correction, a multi-step noise reduction protocol was applied to optimize signal-to-noise ratio while preserving biological dynamics. Rolling ball background subtraction (50-pixel diameter) removed large-scale intensity variations, followed by Gaussian filtering (σ = 2 pixels) to reduce high-frequency noise (Schindelin et al., 2012; Berg et al., 2019). Parameters were empirically optimized to maintain vesicle edge definition while minimizing background fluctuations. Maximum intensity projections of z-planes were then generated to produce high-contrast 2D representations, facilitating robust tracking of vesicular movement over time.

Organelle tracking was performed using the TrackMate algorithm, enhanced with machine learning-based detection and Linear Assignment Problem (LAP) trackers (Tinevez et al., 2017; Simon Youssef et al., 2011). Spot detection employed Laplacian of Gaussian filtering, followed by feature extraction of intensity, size, and contrast, with track linking incorporating gap closing and split/merge handling. Automated tracking results were validated against manual analysis, demonstrating high correlation (r > 0.90) for key movement parameters (Nicolas Chenouard et al., 2014; Yichen Wu et al., 2019). The pipeline extracted multiple movement parameters, including instantaneous and mean velocities (μm/s), maximum speed and acceleration, total displacement, path length, directional persistence, and pause frequency and duration. For chemogenetic experiments involving hM4D(Gi) receptor activation, trafficking parameters were compared within the same axons pre- and post-J60 administration, enabling paired analysis using paired t-tests in GraphPad Prism. Quality control was implemented throughout the pipeline, including semi-automated removal of out-of-focus frames, validation of drift correction via stationary landmarks, monitoring of signal-to-noise ratio across time series, and manual spot checking for tracking accuracy.

#### Image processing, GRAB_NE2h signal analysis, combined with amphisome tracking analysis

Initial image preprocessing was performed in ImageJ-FIJI and Jupyter Lab. Motion correction was applied using the NoRMCorre algorithm with sub-pixel accuracy (Zhou et al., 2018), an essential step for maintaining spatial precision, particularly when tracking small, rapidly moving particles. Noise reduction combined wavelet-based denoising with adaptive thresholding, optimized for neural imaging to preserve biological signal while minimizing background fluctuations (Giovannucci et al., 2019). Organelle tracking was performed using the TrackMate algorithm, enhanced with machine learning-based detection and LAP trackers (Tinevez et al., 2017). This semi-automated approach enabled precise quantification of multiple movement parameters, including instantaneous and mean velocities, displacement vectors, directional persistence, and pause characteristics. Tracking accuracy was validated against manual analysis in subset datasets, confirming the reliability of automated measurements (Chenouard et al., 2014). This approach allows detailed comparison of vesicular behavior across experimental conditions while maintaining consistent measurement criteria. Norepinephrine dynamics were analyzed via GRAB_NE2h fluorescence using specialized ROI selection algorithms (Feng et al., 2019; Feng et al., 2025). Signal processing included adaptive baseline correction (8th percentile F0), photobleaching compensation via exponential fitting, and movement artifact correction through cross-correlation analysis, ensuring signal fidelity for temporal analysis of neurotransmitter dynamics (Patriarchi et al., 2020). To examine peri-event dynamics of the fluorescent biosensor signal, a trigger-based epoch extraction approach was employed. Behavioral and kinematic data were recorded continuously alongside fiber photometry signals sampled at 30 Hz pre averaging. For each recording session, time-stamped events were identified according to criteria defined for the specific experimental question. These event timestamps served as triggers around which neural signal epochs were extracted.

Triggers were defined based on a threshold-crossing criterion applied to a continuous behavioral or kinematic variable “speed, directionality, and displacement”. A valid trigger required: (1) the variable to remain within defined threshold for at least a specified preceding lookback duration, (2) an upward crossing of that threshold at the trigger time point, and (3) sustained activity above the threshold for a minimum post-event duration. To prevent over-counting of closely spaced events, a minimum inter- trigger interval was enforced. Trigger parameters were adjusted per experimental condition to isolate the behavioral event of interest. For each identified trigger, a signal epoch spanning a fixed pre- and post-event window was extracted from the corresponding fluorescence recording. Epochs were only retained if the full window was contained within the recording time range and comprised sufficient data points. For each extracted epoch, the ΔF/F was computed as: ΔF/F (%)=F_0_F−F_0_×100 where F is the raw fluorescence signal an F_0_ is a rolling baseline estimate. The baseline F_0_ was derived through a multi-step procedure: (1) a minimum filter spanning 25% of the epoch duration, (2) followed by a median filter over 50% of that window size, and (3) Savitzky-Golay smoothing (polynomial order 2, window ≈ 30% of the minimum filter size). Feature extraction and event detection were implemented using custom Python scripts, including hierarchical clustering for stop-point identification and wavelet transform analysis for directional changes. Movement patterns were classified via machine learning, with statistical validation through bootstrapping (Pachitariu et al., 2018). Statistical inference used linear mixed-effects models to account for both fixed and random effects, appropriately handling the nested structure of trials within animals (Saxena et al., 2020). Data visualization grand average traces (mean ± SEM) and individual trial overlays were plotted across the full peri-event window. Summary statistics were visualized as box plots with individual trial scatter and bar plots (mean ± SEM), both with and without normalization to the PRE window baseline. Data visualization additionally employed Z-score normalization and hierarchical clustering of temporal patterns to identify population-level structure. A heatmap of the mean ΔF/F trace was generated for each condition.

#### Cell culture, transfection, immunoprecipitation, immunocytochemistry

HEK293T and COS7 cells were maintained under standard culture conditions as described previously (Oelschlegel et al., 2026). For heterologous co-immunoprecipitation (Co-IP) experiments, HEK293T cells were transiently transfected with combinations of EGFP-, mCherry-, and FLAG-tagged constructs using the calcium phosphate precipitation method as previously reported (Oelschlegel et al., 2026). Primary hippocampal neuron cultures were prepared from embryonic day 18 Sprague-Dawley rat embryos of mixed sex as described previously (Grochowska, 2023) and maintained in Neurobasal medium (GIBCO, Thermo Fisher Scientific) supplemented with B27 (GIBCO), L-glutamine (PAA Laboratories, Pasching, Austria).

Neurons were transiently transfected with plasmid DNA using Lipofectamine 2000 (Thermo Fisher Scientific) according to the manufacturer’s instructions. For immunocytochemistry, neurons expressing FLAG-tagged β2-adrenergic receptors were subjected to live labeling to detect surface receptor populations following isoproterenol stimulation to assess colocalization with LC3-positive compartments. Cells were then fixed with 4% paraformaldehyde/4% sucrose for 10 min at room temperature, washed in PBS, and permeabilized with 0.2% Triton X-100 for 10 min. After blocking (2% glycine, 0.2% gelatin, 2% BSA, 50 mM NH₄Cl in PBS, pH 7.4), samples were incubated with secondary antibodies overnight at 4 °C (1:500). Coverslips were mounted using Mowiol 4-88 (Merck). For live-cell imaging, neurons were transfected with SNAP-tagged LC3, labeled with SiR dye, and imaged under basal conditions and following stimulation with 5 µM isoproterenol using a Leica TCS SP8 STED 3X confocal laser scanning system.

#### Antibodies, chemicals, and software

A complete list of antibodies, chemicals, and software used in this study is provided in the Reagents and Tools table.

#### Quantification and statistical analysis

All statistical analyses were performed using GraphPad Prism 9 (GraphPad Software). Sample sizes (n) are reported either in the figure panels or in the corresponding legends. Depending on the study design and data distribution, appropriate parametric tests were applied. Data are presented as mean ± SEM, and statistical significance was defined as \**p* < 0.05, \*\*\**p* < 0.001.

#### Plotting

Scatter plots represent mean ± standard error of the mean (SEM). Plots were generated in FiJi, Python using SciPy, matplotlib, and NumPy packages or GraphPad Prism 9. Figures were assembled and annotated in Adobe Illustrator 2025.

## ACKNOWLEDGMENTS

The authors gratefully acknowledge M. R. Kreutz for providing laboratory facilities and for insightful discussions during the preparation of the manuscript. We also thank U. Thomas, M. Prigge, and E. D. Gundelfinger for suggestions regarding experimental procedures and for their comments on the experimental outcomes. The authors further acknowledge the professional technical assistance of I. Herbert for AAV production, M. Marunde for support with brain sectioning, and C. Boruzki for assistance with Western blotting. We also thank O. Kobler for assistance with image processing, J. Pakan for sharing the cranial window implantation technique, and M. Prigge for sharing the B6.Cg-Dbh^tm3.2(cre)Pjen/J mouse line, the surgical equipment and chemicals, as well as for preparing Animal License 42502-2-1638 LIN. We further acknowledge A. M. Oelschlegel and J. Kaufhold for their assistance with the 42502-2-1638 LIN license amendment.

## Funding

A.K. was supported by the Schram Stiftung, the LIN special project (together with M. Prigge), as well as by grants from the Deutsche Forschungsgemeinschaft (DFG) FOR5228 RP06 (together with M.R. Kreutz) Project-ID 447288260, and FOR5228 RP11N.

## AUTHOR CONTRIBUTIONS

A.K. acquired funding, conceptualized, and supervised the project. A.A.A.A., M.A.-A. and A.K. performed experiments and analysed the data. H.J. assisted with two-photon imaging experiments and provided materials. A.K. wrote the manuscript and prepared the figures with input from A.A.A.A. and M.A.-A. All authors reviewed and approved the final manuscript.

## DECLARATION OF INTERESTS

The authors declare no conflict of interest

## Data and materials availability

Further details and requests for reagents and resources should be directed to the lead contact and will be fulfilled upon reasonable request.

**Table 1.**
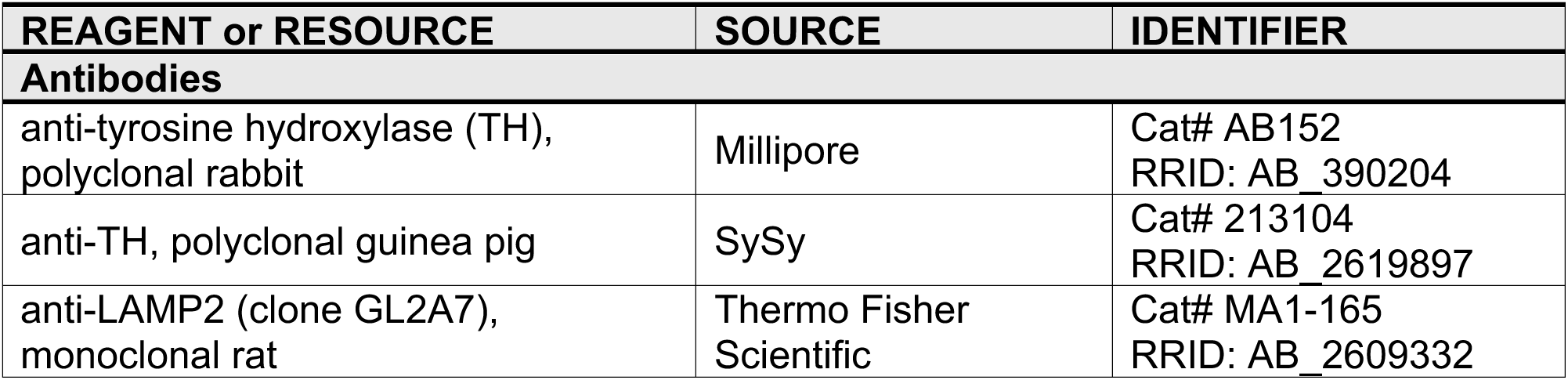
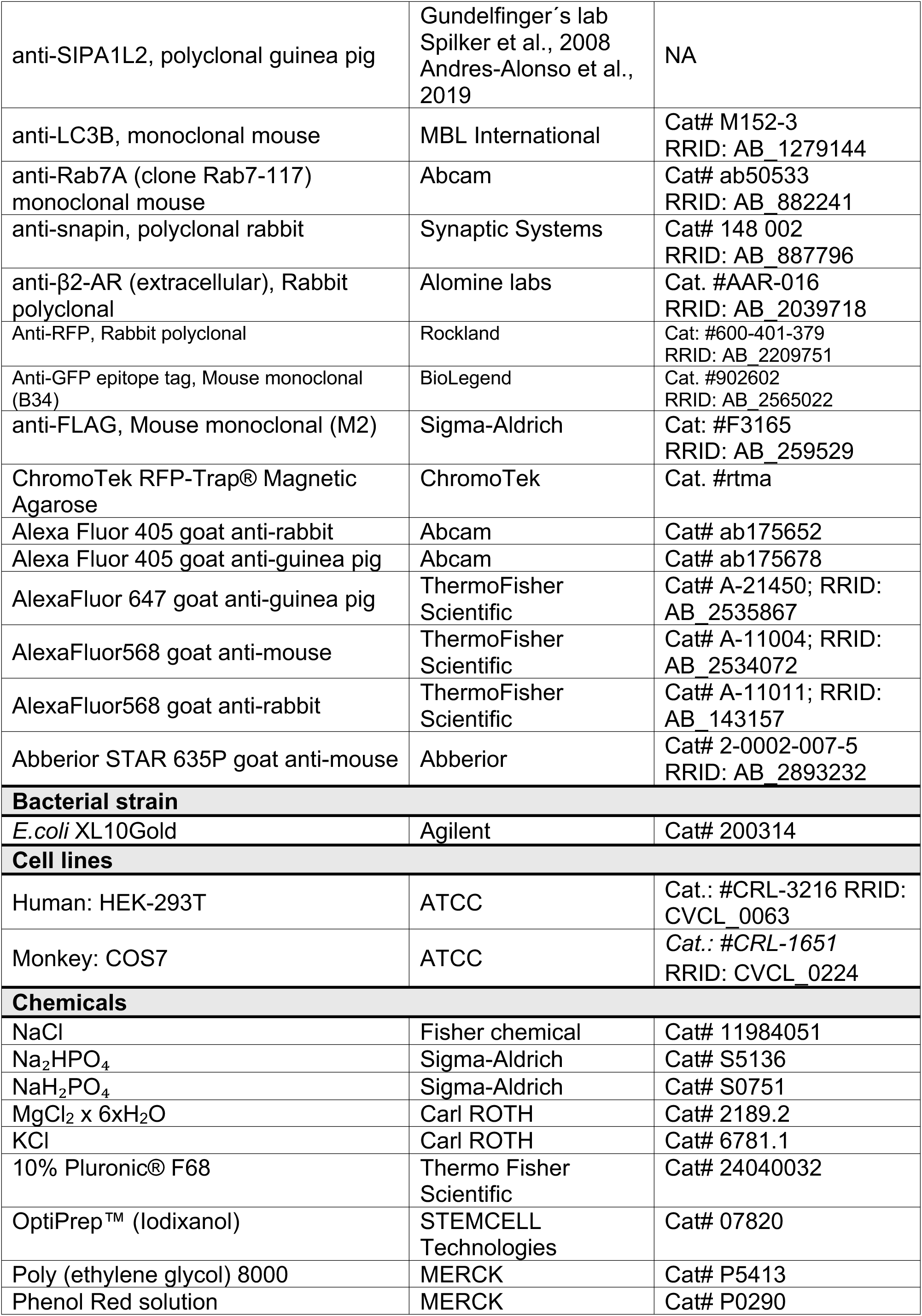
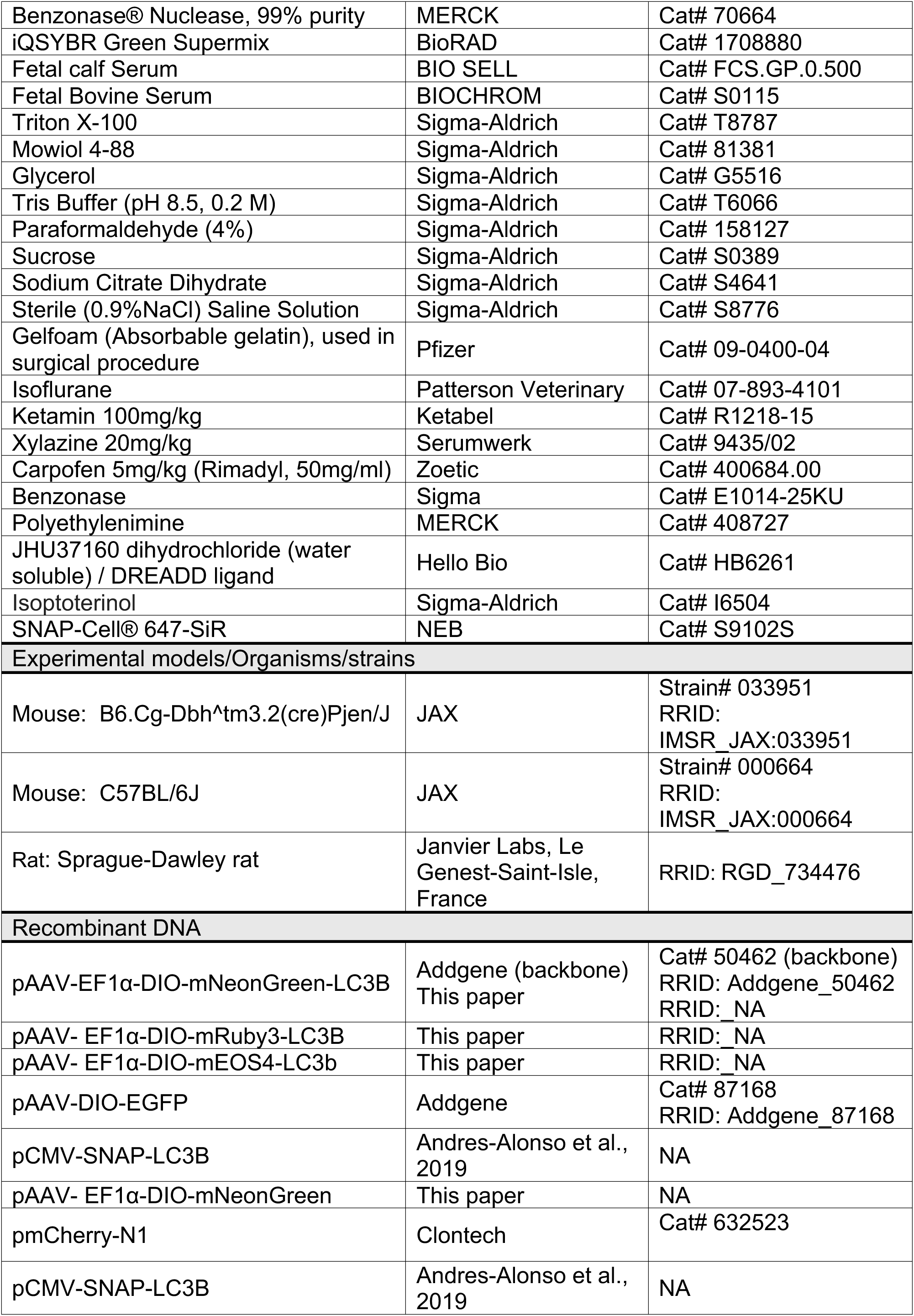
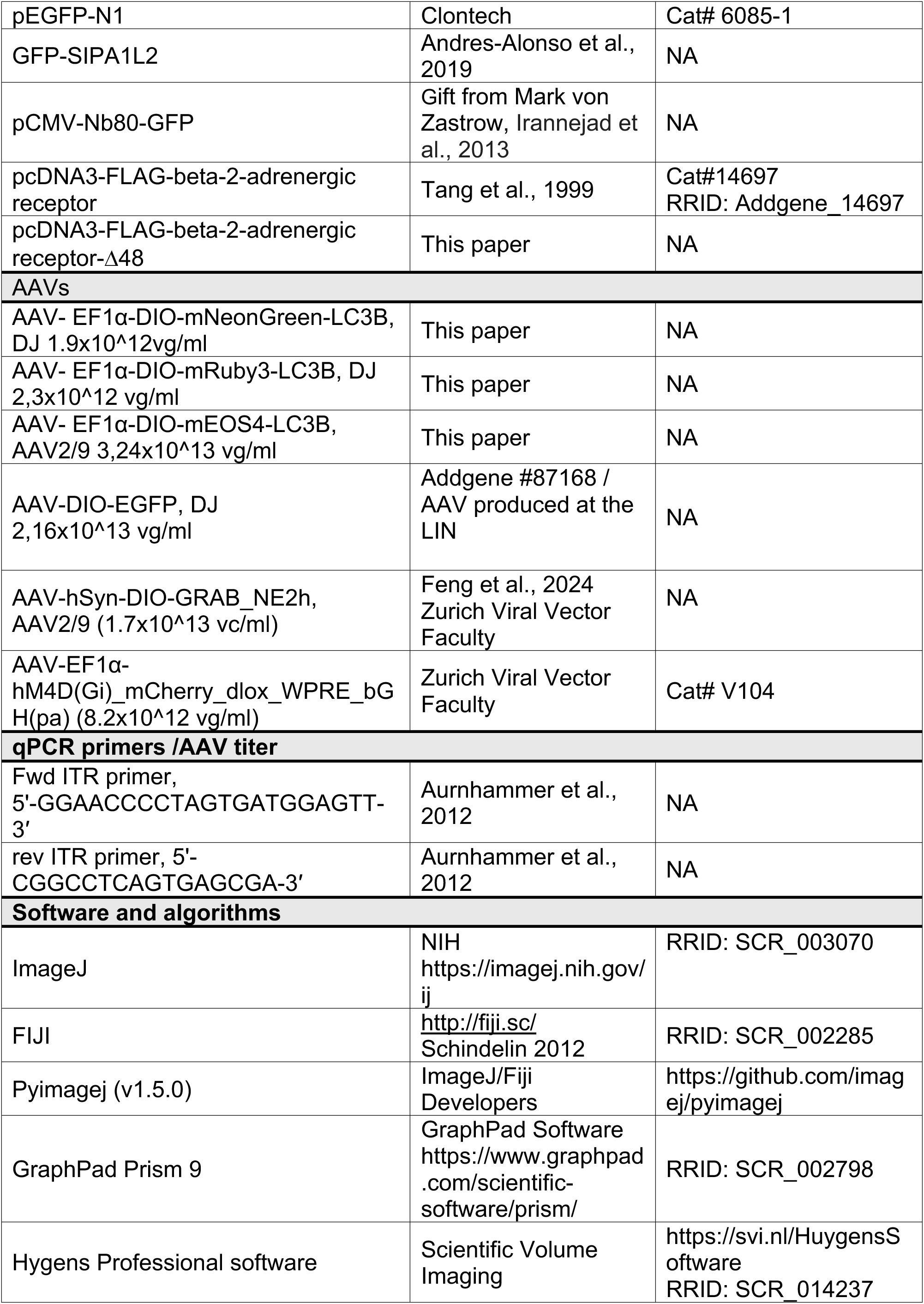
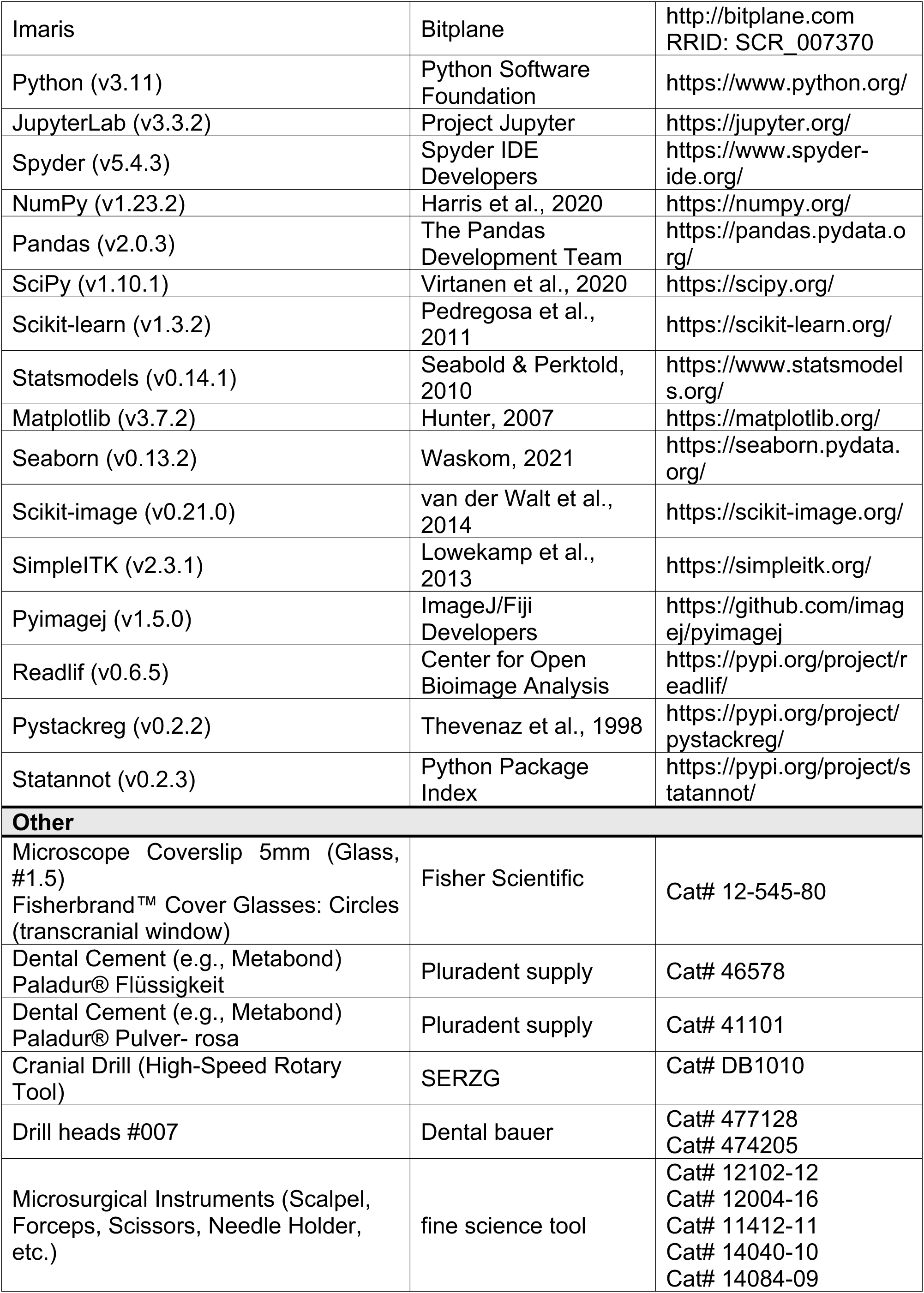
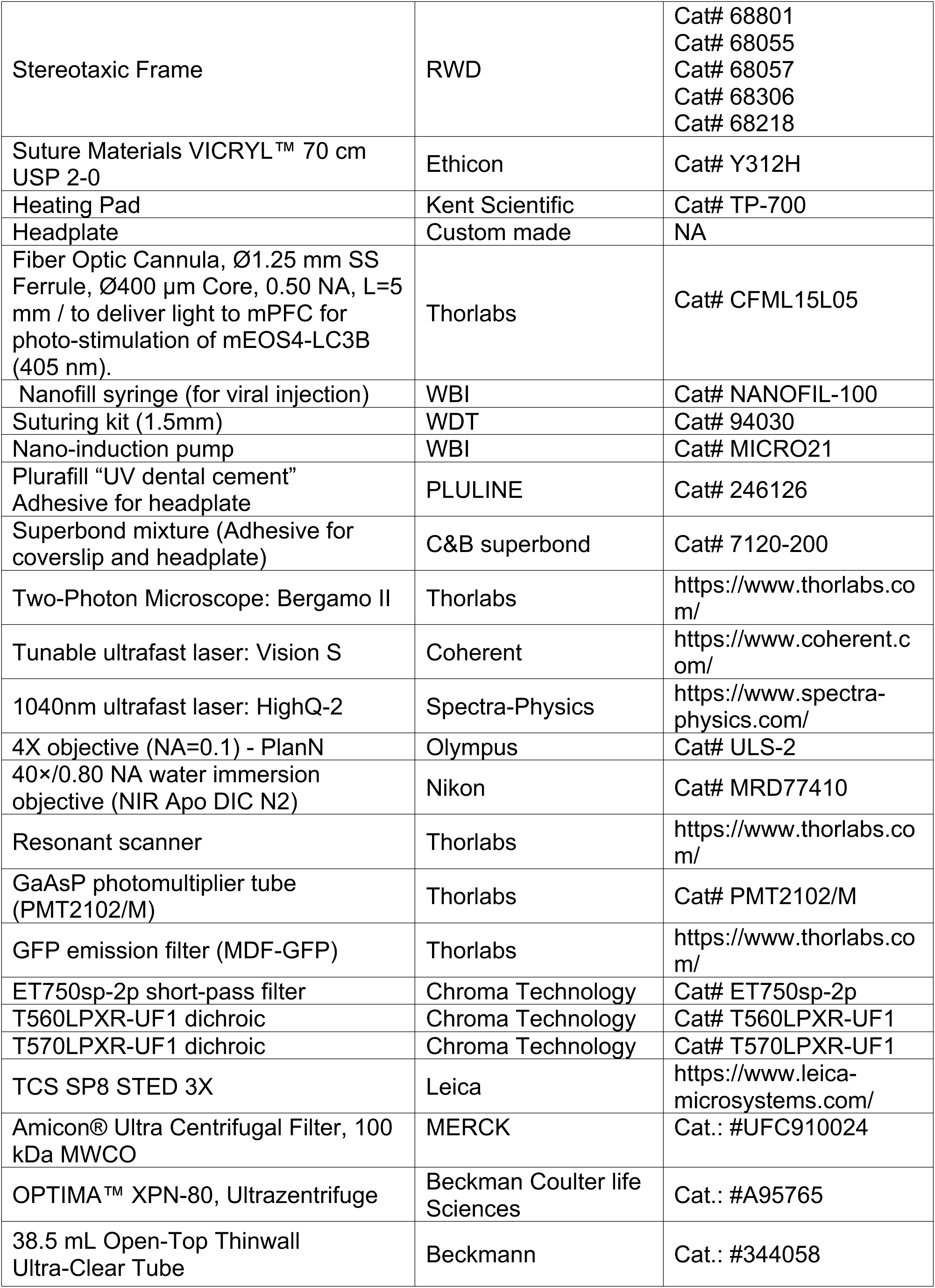
Reagents and Tools.

## SUPPLEMENTAL FIGURES AND LEGENDS

**Figure S1.**
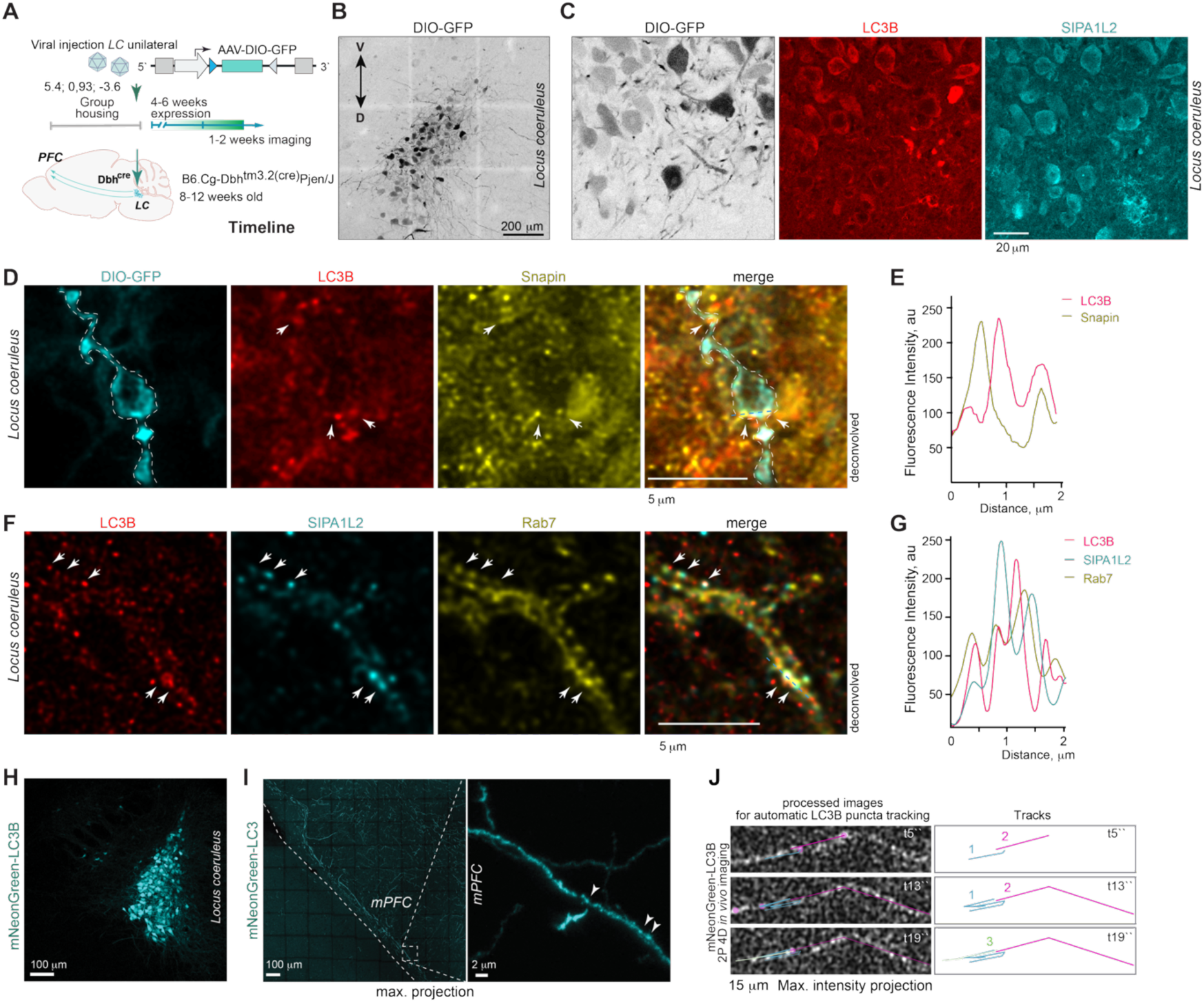
*In vivo*, cell-type-specific labelling of *LC* neurons and axons combined with LC3B-mediated detection of amphisomes. **(A)**. Experimental timeline and AAV-DIO-GFP injections into the *LC* of *Dbh-Cre* mice, expressing Cre recombinase from the endogenous *Dbh* locus. **(B)**. Representative confocal tile-scan image showing specific GFP expression in *LC* neurons. D, dorsal; V, ventral. **(C)**. Representative confocal images of *LC* neurons illustrating LC3B and SIPA1L2 expression. **(D)**. Confocal image of a GFP-labelled *LC* axon with endogenous LC3B and Snapin immunoreactivity, and **(E)** the corresponding line intensity profile. **(F, G)**. Confocal images showing endogenous SIPA1L2, LC3B, and Rab7 co-localized in *LC* neurons. **(H)**. Confocal tile-scan image showing *LC* neurons expressing mNeonGreen-LC3B. **(I)**. Tile scan confocal image of fluorescently labelled *LC* axonal terminals in the *mPFC*, showing dense labelling in layers I and II. Inset highlight LC3B-positive puncta in distal *LC* axons (arrows). **(J).** Semi-automated image analysis of distal *LC* axons showing three distinct tracks over time (1, 2, and 3), representing different trafficking behaviours of LC3B-positive amphisomes *in vivo*.

**Figure S2.**
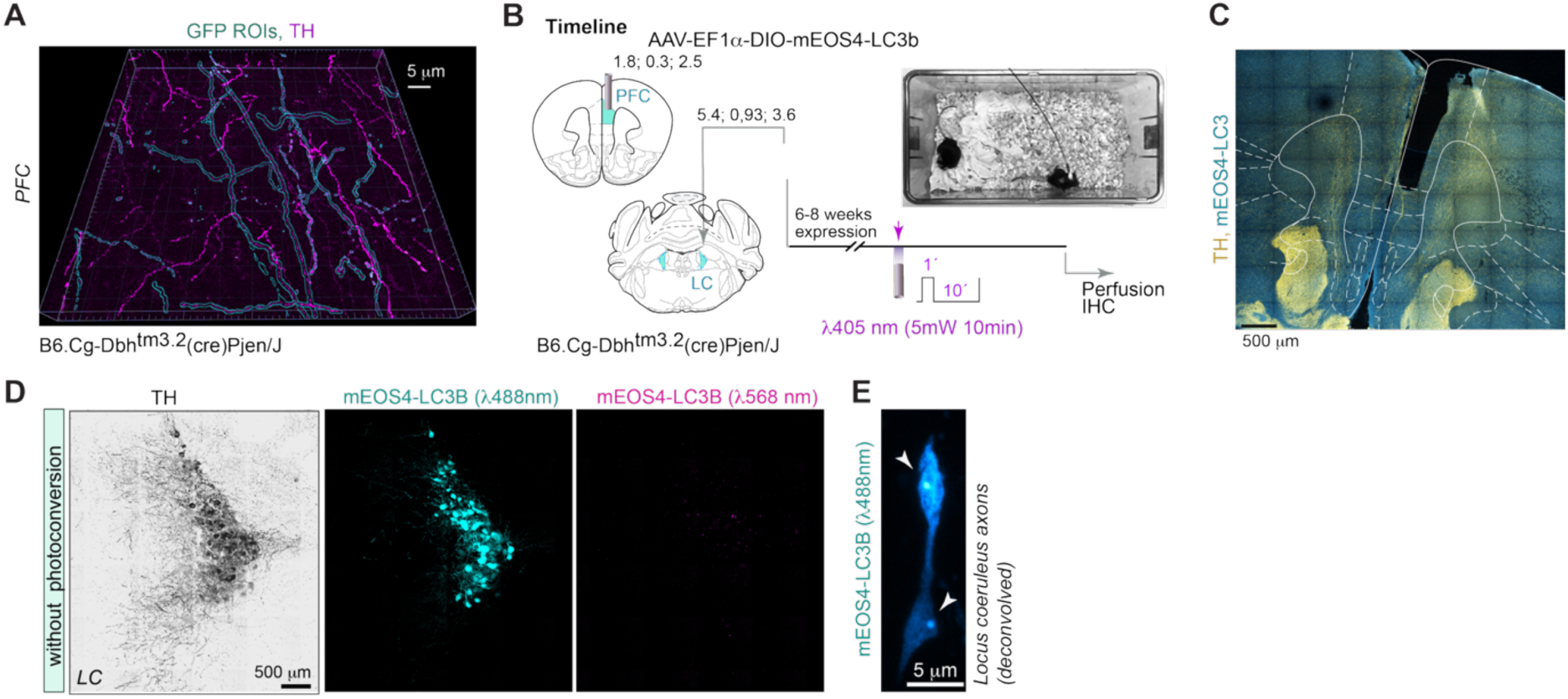
Optical fiber-mediated photoconversion of mEOS4-LC3B in *LC* axons projecting to the *PFC*. **(A)**. GFP-positive axons in the *PFC* used for LAMP2A detection are TH-positive. Colocalization of GFP-labelled LC axons (cyan) with TH immunoreactivity (magenta) confirms *LC*-specific labelling. **(B)**. Experimental timeline: animals were perfused four to six hours after photostimulation, and *LC* sections were immunostained for tyrosine hydroxylase. The photoconverted group underwent a UV-light stimulation protocol consisting of 1-second pulses delivered at 10-second intervals for 10 minutes **(C)**. Representative optical fibre trajectory targeting the *mPFC*. **(D)**. Representative confocal tile-scan image of the *LC* without photostimulation in the *PFC*. **(E)**. Representative confocal image showing mEOS4-LC3B-positive puncta localized within *LC* axonal varicosities, resembling synaptic boutons.

**Figure S3.**
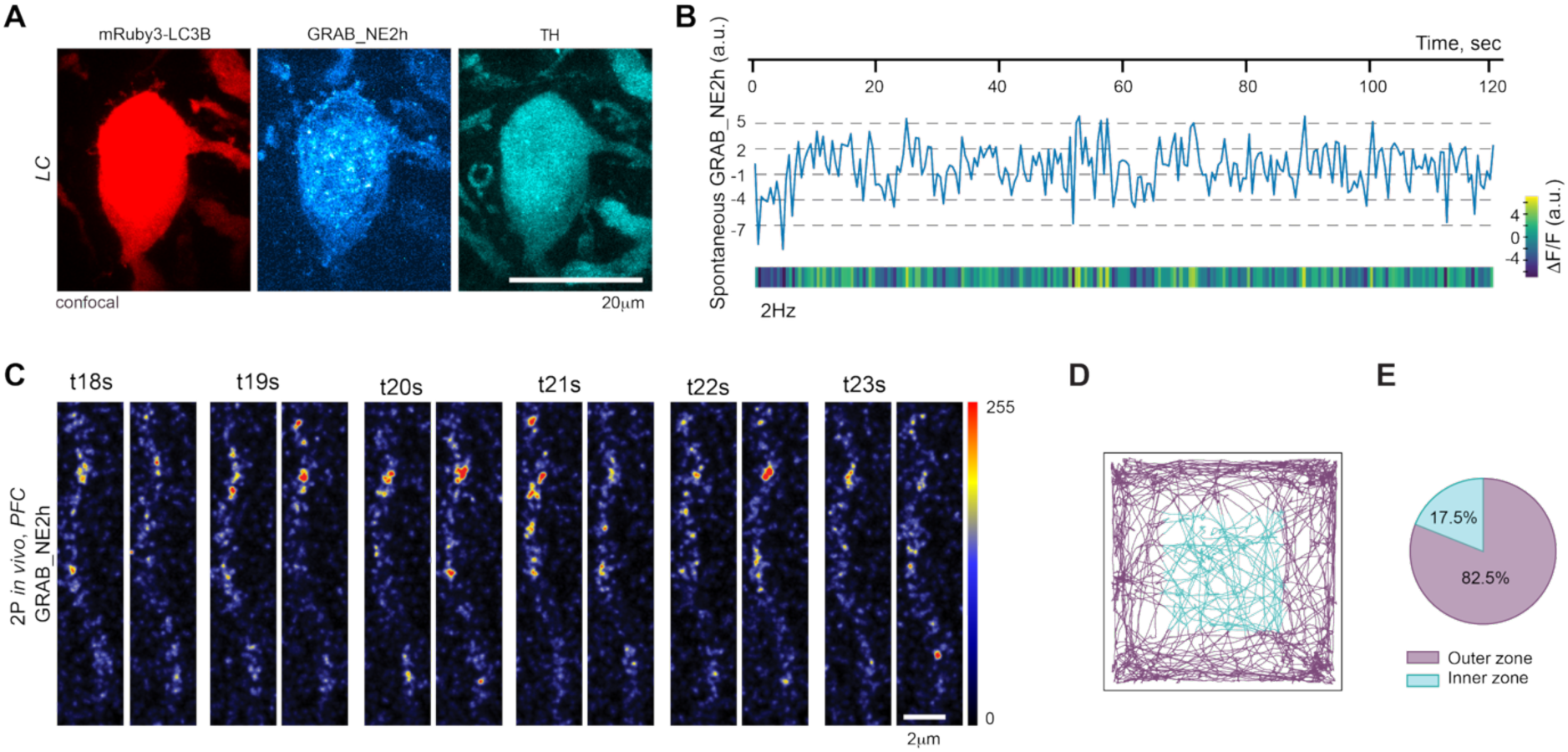
Basal NE dynamics in distal locus coeruleus axons monitored by GRAB_NE2h two-photon imaging *in vivo*. **(A).** Representative confocal image of a *LC* neuron in *Dbh*-Cre mice showing Cre-dependent coexpression of mRuby3-LC3B and GRAB_NE2h. **(B).** Representative trace of spontaneous GRAB_NE2h fluorescence fluctuations recorded by two-photon imaging through a cranial window in distal locus coeruleus noradrenergic axons within the prefrontal cortex. **(C).** Representative two-photon imaging frames illustrating spontaneous fluctuations in GRAB_NE2h fluorescence during basal activity in distal *LC* axons projecting to the *PFC*. **(D)** Visualization of exploratory behavior during photoconversion experiments. Gray-violet traces indicate animal trajectories within the outer zone of the open-field arena, whereas cyan traces denote trajectories within the center zone. **(E)** Pie chart analysis showing the average relative time spent in the outer versus center zones of the open-field arena, indicative of anxiety-like behavior.

**Figure S4.**
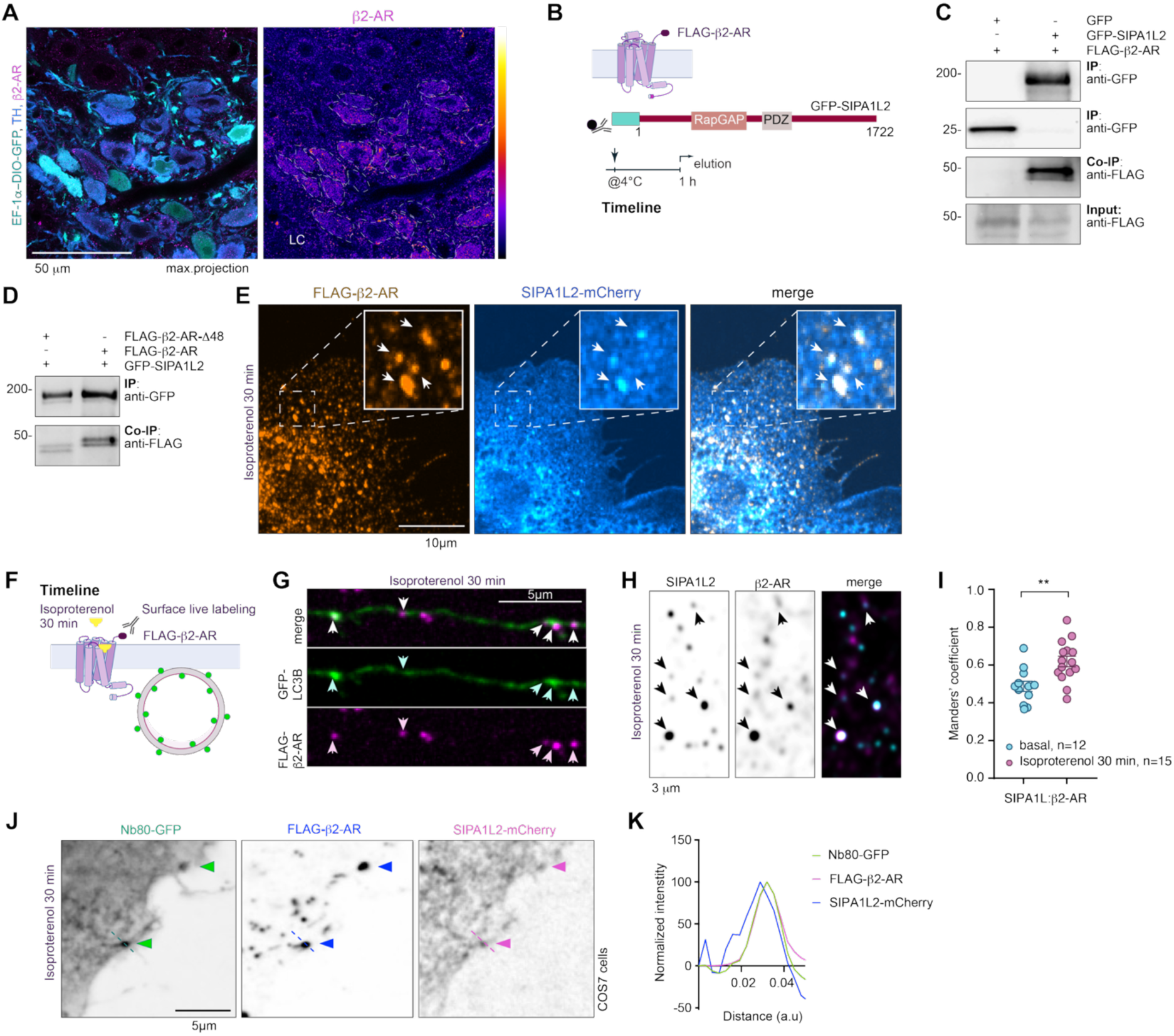
β2-adrenergic receptor activation promotes recruitment of SIPA1L2-positive amphisomes. **(A)**. Representative confocal image showing Cre-dependent GFP expression in TH-positive *LC* neurons and somatic β2-AR immunoreactivity. **(B)**. Schematic of the β2-AR-SIPA1L2 co-immunoprecipitation experiment. **(C)**. Immunoblot showing co-immunoprecipitation of β2-AR with SIPA1L2. **(D)**. SIPA1L2 association with β2-AR is mediated by the receptor C-terminus containing the PDZ-binding motif. **(E)**. Representative confocal image showing co-localization of β2-AR and SIPA1L2 within vesicular compartments in COS7 cells. **(F)**. Schematic of the experimental timeline for live labeling of FLAG-tagged β2-AR in primary hippocampal neurons. **(G)**. Representative confocal images showing proximity of LC3B-positive puncta to surface-labeled β2-AR in axons of hippocampal neurons treated with isoproterenol (5 μM). **(H, I)**. Isoproterenol treatment (5 μM) enhances co-localization of β2-AR and SIPA1L2 in cultured neurons, consistent with recruitment of SIPA1L2-positive vesicles to activated β2-AR. **(J, K)**. SIPA1L2, β2-AR, and the nanobody-based conformational biosensor Nb80 co-localize in COS7 cells following isoproterenol treatment. Nb80-GFP selectively labels activated β2-AR. Relative intensities within the indicated region are shown in the line profile (dashed line in J).

**Figure S5.**
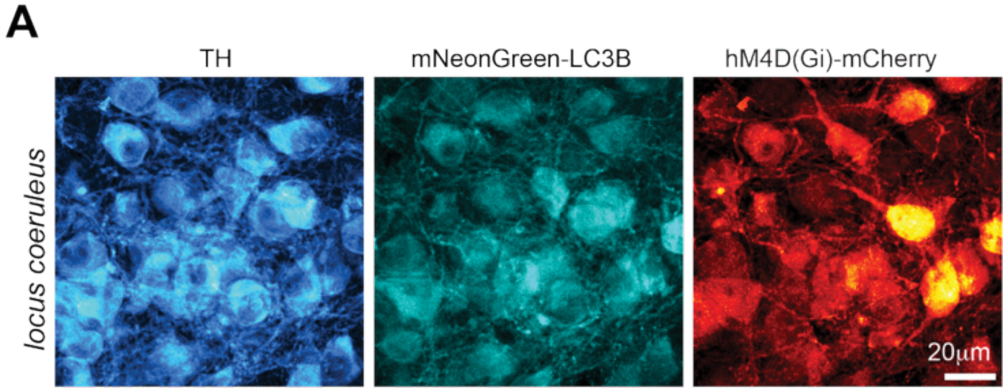
Co-expression of mNeonGreen-LC3B and hM4D(Gi)-mCherry in *LC* neurons. **(A)** Histological validation following co-injection of AAV-DIO-mNeonGreen-LC3B and AAV-hSyn-hM4D(Gi)-mCherry into Dbh-Cre animals. Representative confocal images show co-expression of mNeonGreen-LC3B and hM4D(Gi)-mCherry in TH-positive *LC* neurons. Expression of hM4D(Gi)-mCherry driven by the hSyn promoter was neuron-specific but not restricted to locus coeruleus neurons.

## Notes

### Competing Interest Statement

The authors have declared no competing interest.

### Summary of Updates

The section on β2-adrenergic receptor signaling has been revised to clarify the molecular mechanism underlying amphisome confinement. Most figures have been updated. The author list and supplementary materials have also been updated.

